# N-terminal Chirality and Sequence Variations Modulate the Conformational Landscape of Amyloid-beta 42^†^

**DOI:** 10.64898/2026.03.19.713039

**Authors:** Qiang Zhu, Haibo Yu

## Abstract

Amyloid beta (A*β*), one of the hallmark proteins of Alzheimer’s Disease (AD), aggregates into plaques that are strongly linked to cognitive decline and neuronal death. Reducing its aggregation propensity may provide a strategy to slow the progression of AD. While chirality modulation has emerged as an innovative approach to disrupt this process, research has primarily focused on alterations at the *Cα* position, often overlooking the impact of the second chiral center, such as the *Cβ* atom of Threonine. Furthermore, the underlying mechanisms governing these chiral effects remain elusive. Given the intrinsically disordered nature of the A*β* peptide, we employed temperature-replica exchange molecular dynamics (T-REMD) simulations to explore its rugged conformational landscape. We considered sequence mutations (A2T, A2V), N-terminal chirality inversion of the first six residues (A2V_1−6D_ and WT_1−6D_), and alteration of the second chiral center (C*β*) of Threonine (A2T_*Cβ*_). By analyzing the effect size and population change induced by these mutations and chiral modulation, we concluded that the modulation at the N-termini is not confined locally but also exerts specific effects on the central hydrophobic core (CHC) region. Inspection of their free energy landscape and representative structures reveals that the protective or pathogenic effects of these variants correlate with their similarity to the wild type (WT) ensemble. Beyond these static thermodynamics analyses, a direct connection to phase transitions was made by estimating heat capacity as a function of temperature. Both analyses predict that A2T_C*β*_ may exert a pathogenic effect, in contrast to the protective nature of A2T. These findings offer a deeper understanding of the effects of site-specific mutations and chirality and shed light on the development of advanced therapeutic strategies for AD.

## 1 Introduction

With the aging global population, Alzheimer’s Disease (AD), a major form of neurodegenerative disease, is becoming an increasingly significant public health burden, and the number of individuals diagnosed is projected to reach 13.8 million by 2060. ^1,2^ The aggregation of amyloid beta (A*β*) peptide, produced through the cleavage of amyloid precursor protein (APP), ^3,4^ into mature fibrils and oligomers is recognized as a hallmark of AD pathology.

Among A*β* peptides, the two predominant isoforms are A*β*40 and A*β* 42. ^5^ They share a common structural architecture, comprising an N-terminal region (residues 1-14) and hydrophobic interaction sites, namely the central hydrophobic core (CHC, residues 17–21) and the hydrophobic C-terminal region (residues 30–36), respectively. A*β*42 differs from A*β*40 by two additional residues at the C-terminus. Compared with A*β*40, the two residues render the variant A*β*42 more prone to aggregation and neurotoxicity. ^6–8^

Significant efforts have examined the effects of sequence modification, ^9–11^ length, ^12–14^ and external factors (such as small molecules, ^15^ receptors, ^16^ metal ions, ^17^ and cell membranes ^18^) through both experimental and computational approaches. ^19,20^ For instance, a causative mutation A2V (a substitution of alanine with valine at site 2 of A*β*42) has been shown to increase the risk of AD in the homozygous condition. ^21^ Conversely, the substitution of alanine with threonine A2T exerts a protective effect against AD and A*β* assembly. ^22^ Furthermore, mutations adjacent to CHC, such as E22G ^23^ and E22K ^24^ variants, significantly accelerate aggregation by altering the electrostatic environment and enhancing hydrophobic packing. Beyond sequence mutations, chirality modulation has emerged as an innovative strategy to probe and control amyloid aggregation, ^25–28^ driving growing interest beyond sequence-level approaches. The literature reports that chiral alteration of the first six residues in WT (WT_1−6D_) and A2V (A2V_1−6D_) peptides protects cells from the toxicity induced by A*β* fibrillization. ^29^ Parallel to these N-terminal modifications, even single D-amino acid chirality alterations at critical internal positions are sufficient to incur substantial effects. For instance, the chiral editing of K16 and V24 has been shown to effectively inhibit fibril formation. ^11^ Interestingly, this sensitivity to chiral inversion is not uniform across all amyloid species. Through an unbiased D-amino acid substitution strategy, researchers have disclosed that the aggregation landscape of A*β*42 is significantly more sensitive to such chiral perturbations than that of its shorter counterpart, A*β*40. ^27^

To account for the distinct effects of different variants, a range of mechanisms has been proposed based on both experimental assays and computational simulations to explain their divergent aggregation propensities. These explanations predominantly focus on global parameters such as side-chain hydrophobicity, ^30^ structural ensemble ^31–35^ and stability ^36^, steric bulk, ^37^ and solvation free energy ^38^. However, despite the comprehensive nature of these studies, a systematic investigation into the role of side-chain chirality remains limited. Crucially, the fact that Threonine (Thr) possesses a second chiral center at the C*β* position ^39^ (Figure 1) has been largely overlooked in the context of A*β* aggregation modulation. Considering the limited success of clinical trials ^40^ targeting the CHC regions, there is a growing realization that the N-terminal region may hold the key to understanding the early-stage modulation of A*β* aggregation. ^41,42^ However, the unstructured nature of the N-terminal region ^43,44^ poses a significant challenge for conventional experimental techniques, such as X-ray crystallography or cryo-electron microscopy, which often struggle to resolve such highly flexible and transient structural ensembles. Consequently, the molecular-level details of how N-terminal modifications, particularly side-chain chirality, govern the early stages of A*β* self-assembly remain elusive.

**Fig. 1.**
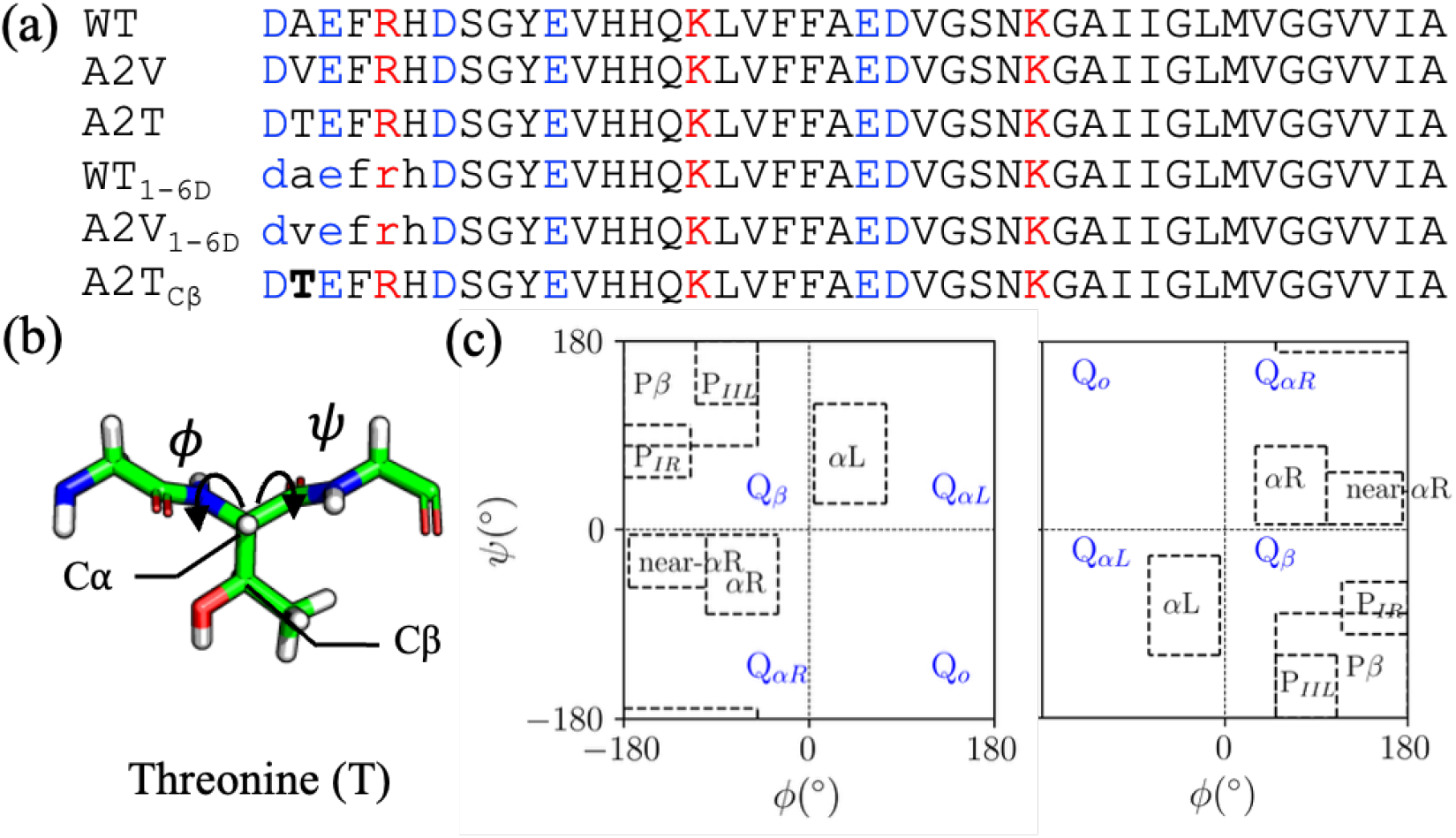
Illustration of the systems investigated in this study. (a) Residue sequences of the six variants, namely, WT, A2V, A2T, WT_1−6D_, A2V_1−6D_, and A2T_C*β*_, respectively. Residues with L-chirality are shown in uppercase, while those in which the C*α* chirality has been inverted are shown in lowercase. Alterations in C*β* chirality are indicated in bold. Positively charged and negatively charged residues are colored in red and blue, respectively; (b) Structural representation of dihedral *ψ, ϕ*, and the position of C*α* and C*β* of Threonine (T); (c) Defined quadrants and mapped regions for both the L-and D-amino acids. For L-amino acid, their conformational regions are defined as below: *α*R (*ϕ* ∈ [−100°, −30°], *ψ* ∈ [−80°, −5°]), near-*α*R (*ϕ* ∈ [−175°, −100°], *ψ* ∈ [−55°, −5°]), *α*L (*ϕ* ∈ [5°, 75 °], *ψ* ∈ [25°, 120 °]), P*β* (*ψ* ∈ [−180°, −50°], *ψ* ∈ [80°, −170°]), P_IIL_ (*ϕ* ∈ [−110°, −50°], *ψ* ∈ [120°, 180°]), P_IR_ (*ϕ* ∈ [−180°, −115°], *ψ* ∈ [50°, 100°]). For D-amino acids, all regions correspond to the inversion of the L-amino acid map through the origin. These regions definitions are taken from the literature^39^ directly.

Complementary to experimental techniques, computational studies provide invaluable insights into the diverse conformational ensembles that are often experimentally inaccessible. While cytotoxicity is primarily associated with soluble oligomeric intermediates formed during early-stage aggregation, ^45–47^ a detailed characterization of the A*β* monomer is merited for deciphering polymerization pathways and designing strategies to inhibit oligomerization at its inception. ^33,35,48,49^ In this context, we used all-atom molecular dynamics simulation and focused on A*β* 42, which exhibits heightened neurotoxicity and is central to AD pathology. We focused particularly on the N-terminal region. We modeled the wild-type (WT) peptide alongside the A2T and A2V variants, specifically incorporating chirality alterations at the C*β* position of Threonine. Since A*β* belongs to the class of intrinsically disordered proteins (IDPs) ^50^ characterized by a rugged free energy landscape, temperature-replica exchange molecular dynamics (T-REMD) simulations ^51^ were employed to achieve robust conformational sampling. Through comprehensive analysis, we revealed that the modulatory effects of the N-terminal region are not locally confined; instead, they propagate to CHC regions.

Furthermore, we demonstrated that backbone chirality (C*α*), including the often-ignored second chiral center (C*β*) of Threonine, reshapes the conformational landscape. These findings underscore the critical role of chiral configuration in the aggregation process, delivering molecular insights into the development of targeted therapeutic interventions.

## 2 Computational Details

The initial structures of A*β*42 and its variants were generated in an extended conformation using the *tleap* module ^52^, rather than using compact NMR-derived structures under non-physiological conditions, as noted previously ^9,53^. The initial structures of the chirality-inverted variants, namely, WT_1−6D_, A2V_1−6D_, and A2T_C*β*_, were generated with the help of ChimeraX ^54^. Here, we focus on the conformational landscape rather than their kinetics. Based on previous studies demonstrating the reliable performance of Amber99SB ^55^ in simulations of the A*β*42 systems ^34,56^, we adopted the Amber99SB force field with the generalized Born (GB) implicit solvation model, consistent with established protocols. ^34^ A surface tension coefficient was set to be 0.005 *kcal/mol/*Å^2^ to incorporate the nonpolar solvation effects. A salt concentration was set to be 0.2 M.

14 temperature replicas ranging from 287 to 450 *K* following an exponential distribution (287.00, 297.10, 307.56, 318.39, 329.60, 341.20, 353.21, 365.65, 378.52, 391.84, 405.64, 419.92, 434.70, 450.00 *K*) were employed here as reported previously. ^53^ The temperature was manipulated using Langevin dynamics with the collision frequency set to be 1 *ps*^−1^. The exchange between replicas was attempted at intervals of 1 *ps*. The exchange probability between any paired replicas (*Prob*_*i*→*j*_) was maintained between 18 % and 22 % through the Metropolis criterion to preserve thermodynamic ensemble, as detailed in Eq. 1. Such a range has been demonstrated to be optimal for T-REMD, where a low exchange probability will contaminate the sampling efficiency by hindering temperature-space diffusion, while a high rate prevent structures from sufficiently relaxing at the desired temperature. ^53,57^ All bonds involving hydrogen atoms were constrained with the SHAKE ^58^ algorithm and the integration time step was set to 2 *f s*.

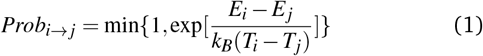

where *Prob*_*i*→ *j*_ denotes the exchange probability between replicas *i* and *j, E* and *T* of different subscript *i* and *j* are their corresponding energies and temperature, respectively.

Each temperature replica of the variants was simulated for 0.8 *µs*, except for A2T, WT_1−6D_, A2V_1−6D_, and A2T_C*β*_, which was run for 1.2 *µs*. A detailed summary of the systems investigated in this study can be found in Figure 1(a) and Table S1. The simulation timescales required for convergence in these T-REMD runs align well with those reported by several other groups, ^31,44,53^ and were necessary to mitigate potential force-field-specific artifacts as previously documented. ^53^ All simulations were conducted using AMBER package suite ^59^ and analyzed using the *cpptraj* module ^52^ and in-house python scripts with trajectories processed with package *mdtraj* ^60^. Dimensional reduction and clustering were performed using *scikit-learn*. ^61^ All scripts used in this work are available at https://github.com/QZResearch/Abeta42

## 3 Results and Discussion

### 3.1 Convergence and Performance Estimation

Demonstrating convergence and validating the adopted force field are prerequisites for the analyses below. To assess convergence, we examined the distribution of radius of gyration (*R*_*g*_) and the population of secondary structure propensity at specific residues. With the sufficient simulation time, the distribution profile of *R*_*g*_ should be similar at two separate time intervals. For secondary structure propensity, its accumulated population should level off as increasing the simulation time, reflecting frequent transitions between conformational states. Here, we focused exclusively on the replica simulated at 297 K, as this temperature is physiologically relevant.

As demonstrated in Figure S1, the accumulated secondary structure propensity reaches a plateau around 0.4 *µs* in most cases. For the *R*_*g*_ distribution analysis, the first several hundred nanoseconds of simulation were discarded for equilibrium. The *R*_*g*_ distributions at two different time intervals agree well with each other (Figure S2), further supporting convergence. We further estimated the residue-wise secondary-structure populations across 14 temperature replicas. As shown in Figure S3, at lower temperatures, individual residues exhibit distinct propensities for *β*-sheet and *α*-helical conformations. The *β*-sheet profile is consistent with previously reported data. ^53^ However, as the temperature increases, these structured propensities diminish, while the population of bend and turn conformations rises and becomes evenly distributed. It further demonstrates the efficiency of the protocol we employed here. Subsequent analyses were performed using the final 400 *ns* of the trajectories for A2T, WT_1−6D_, A2T_C*β*_, and A2V_1−6D_, and the final 320 *ns* for WT and A2V, respectively.

The performance of the model was validated against experimental chemical shifts obtained from two independent datasets ^62,63^ and values predicted by LEGOLAS ^64^ based on the simulated trajectories. For these comparisons, the replica simulated at 287 *K* was utilised, as this temperature most closely approximates the experimental conditions reported in determining chemical shift. As shown in Figure S4, strong correlation was observed for C*α* and C*β* (R^2^ *>* 0.9). However, the weaker correlation observed for amide N and H*α* shifts likely reflects their strong dependence on hydrogen-bond geometry and local electrostatic environments. This discrepancy mainly arises from the limitations of the implicit solvent model, as the absence of explicit water molecules may hinder the accurate representation of transient hydrogen-bonding interactions. In conclusion, these results demonstrate that the ensemble sampling is adequate and the parameters chosen are appropriate for subsequent analyses.

### 3.2 Conformational Sampling of *ϕ***/***ψ* Space

A detailed examination of residue-wise conformational preferences was performed by inspecting the conformational sampling of *ϕ* /*ψ* space. This space was partitioned into four quadrants of the Ramachandran plot and further divided into six specific conformational regions, namely, *α*R, near-*α*R, *α*L, P*β*, P_IIL_, P_IR_. The precise definitions for these regions are provided in Figure 1(c), following Ref. ^39^. Raw data displaying the Ramachandran plots for each residue of the variants are provided in the Supporting Information (Figures S5–S10).

To estimate whether the conformational propensity changes are significant compared to WT, we employed a previously reported approach for quantifying the similarity between L- and D-amino acid ^39^. For D-amino acids, these regions were simply reflected through the origin, where *ϕ* and *ψ* equal 0. Two properties, namely, the effect size (*E*) and conformation population changes (|Δ%|), were used. Their definitions are detailed as below:

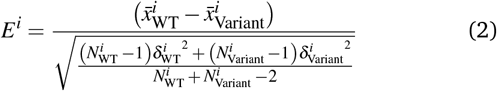

where *N* denotes the number of data points collected within a certain region, 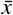 represents the mean value of *ψ* and *ϕ, δ* is for the standard deviations of these data. Subscripts WT and Variants denote wild type and distinct variants investigated in this work, respectively. Superscript *i* represents the *i*_*th*_ residue we investigated.

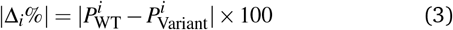

where *P* denotes the population of certain conformational regions. The superscripts and subscripts used here follow the conventions defined in Eq. 2.

As expected, distinct effect sizes across *ϕ* /*ψ* space and significant population shifts were observed within the first six residues, where the mutations and chiral modulations were introduced (Figure S11). Using an effect size threshold of 0.5, which is considered a substantive difference ^39^, predominant effect sizes were observed in the *α*R and *α*L regions for variants WT_1−6D_ and A2V_1−6D_ compared to other conformational regions. In addition, the effects exerted by single mutations and chiral modulations were consistently observed to be indirect across all investigated sites. For instance, although modifications in the A2V, A2T, and A2T_*Cβ*_ variants were introduced solely at site 2, significant differences were observed at distant residues as well. In the A2T variant, the maximum perturbation in the *ϕ* dihedral angle within the near-*α*R region was observed at site 3, while for the *α*_L_ region, the peak effect of *ϕ* and *ψ* shifted further to site 6. In the A2V variant, similar trend was observed and a second peak was detected in the *ψ* dihedral within the *α*_L_ region.

When the effect sizes and population changes at residues excluding the initial six positions were examined, more distinct trends were observed. As illustrated in Figure 2, the effect sizes within the *α*R region were found to be less significant than those in other conformational regions. Its conformational population changes were primarily localized to the residues adjacent to the modification site. Across these variants, the population changes induced by WT_1−6D_ and A2V_1−6D_ are more pronounced than those observed in other variants. This heightened sensitivity in these structural boundaries may reflect the backbone tension and steric strain triggered by the incorporation of D-amino acids that have been observed in using D-enantiomeric peptides for disassembling tau fibrils. ^28^ In addition, a peak was observed between residue 30 and 36.

**Fig. 2.**
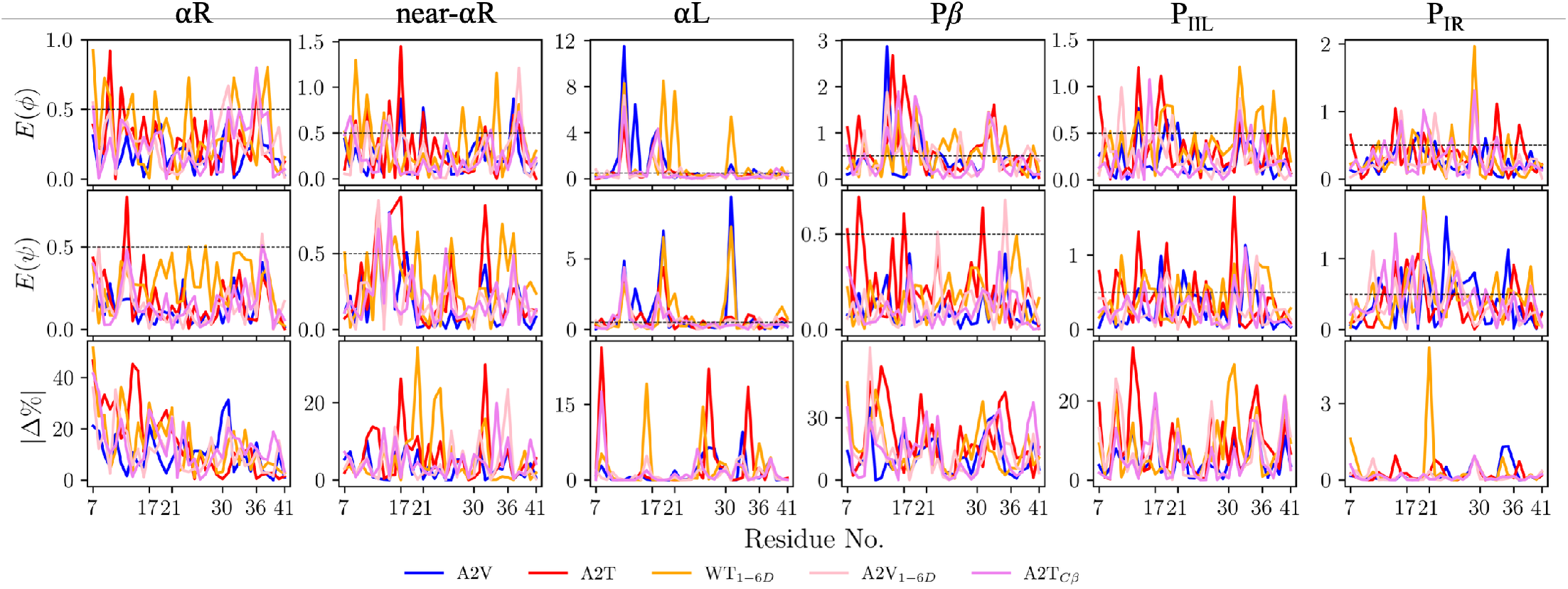
Effect size analyses of both *ϕ* and *ψ* and population change percentage |Δ%| sampled among different variants and as a function of residue index. The effect size threshold of 0.5 is indicated by a dashed line.

In contrast, significantly larger effect sizes of *ϕ* /*ψ* spaces were identified for the *α*L, P_IIL_, and P_IR_. Most interestingly, among these conformational regions, two distinct peaks were identified and localized within or adjacent to residues 17–21 and 30–36. These two regions fall into the CHC region which have been previously demonstrated to be responsible for the fibril formation. ^65,66^ This observation establishes a connection between the well-known aggregation-prone CHC regions and the intrinsic disordered N-terminus, further supporting the N-terminal domain as a promising target for therapeutic intervention. ^41^

### 3.3 Changes in Free Energy Landscape

By projecting the high-dimensional conformational space onto a basis of backbone dihedrals, the primary degrees of freedom governing proteins’ secondary structures are effectively captured. ^67^ This approach is particularly advantageous for capturing metastable states and rare transitions, providing a robust representation of the rugged free energy landscape. For the purpose of visualization, the first two principal components (PCs) are displayed in Figure 3. We observed that even single amino-acid modifications result in distinct free-energy shapes, reflecting substantial alterations in the underlying conformational equilibria of the A*β* variants. Furthermore, the presence of multiple basins is identified within these landscapes. This aligns with the inherent conformational heterogeneity of intrinsically disordered proteins (IDPs).

**Fig. 3.**
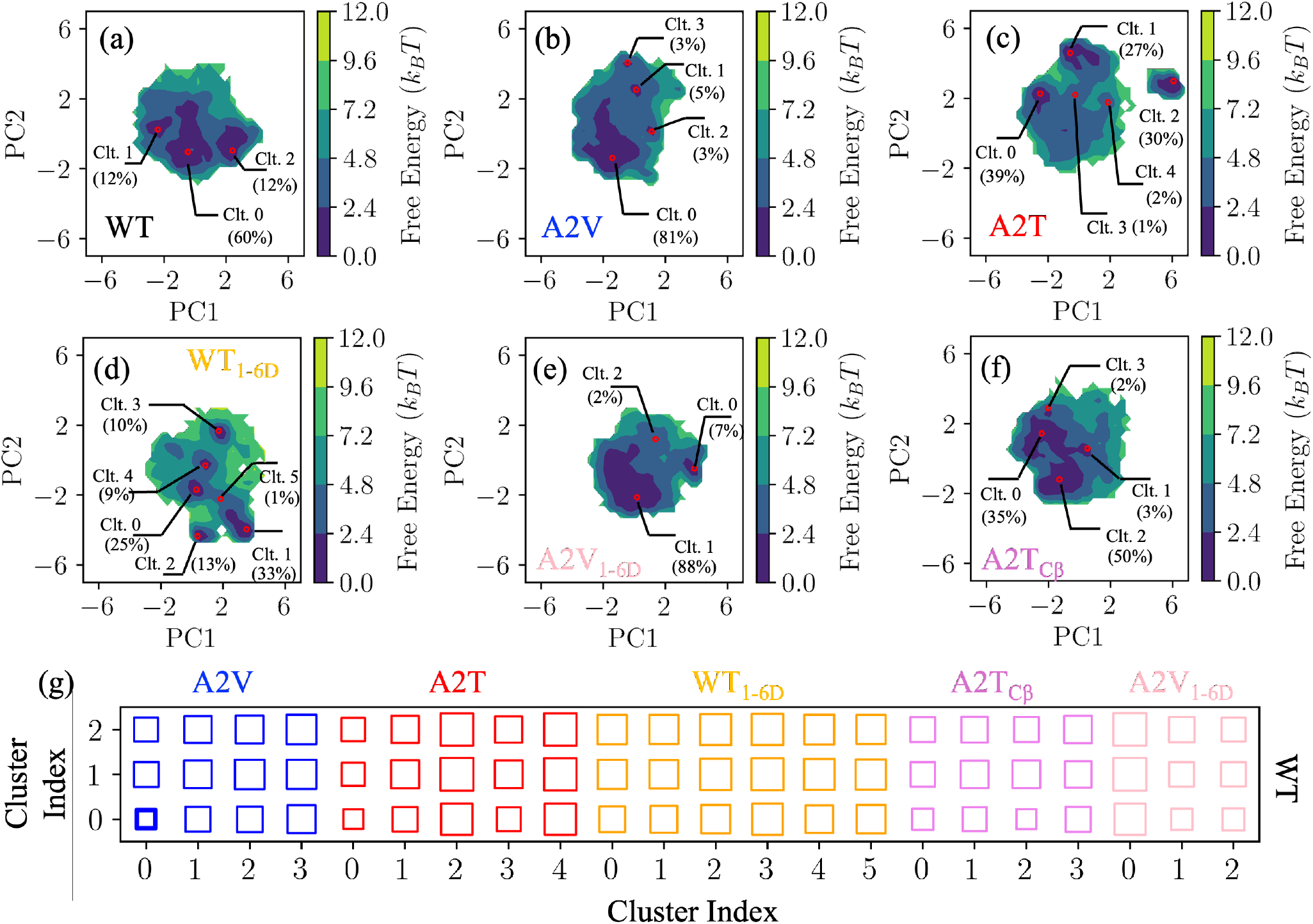
Energy landscape of six variants mapped onto the two principle component (PC) coordinates: (a) WT; (b) A2V; (c) A2T; (d) WT_1−6D_; (e) A2V_1−6D_; and (f) A2T_C*β*_. Their representative coordinates are represented using hollow red circle, together shown are their index label and corresponding populations noted in parenthesis. The detailed clustering results can be found in supporting information (Figure S13). On the bottom panel (g) shows the similarity scored between the representative structures of wild type and other variants, incorporating the writhe features for segment length (*l*) 1 and 5. A lower score indicates a higher degree of structural similarity; specifically, differences smaller than 3.5 are highlighted in bold to indicate closely related conformational states.

A density-based clustering method (DBSCAN, density-based spatial clustering of applications with noise) ^68^ was used here to smooth the landscape and identify representative basins. Detailed procedures for key parameters selection, along with the identification of distinct clusters through color-coding, are provided in the Supporting Information (cf. Section S5). For clarity, only the representative structure of each cluster is labeled on the mapped landscape, as determined by Eq. S2.

As shown in Figure 3(a), three local minima were identified on the free energy landscape of WT, with Cluster 0 being the most dominant at 60 %, followed by Cluster 2 (18 %) and Cluster 1 (12 %). In the case of A2V, four minima were detected, with Cluster 0 accounting for 81 % of the population. (Figure 3(b)) While the dominant structures of both WT and A2V occupy a similar region of the free energy landscape, their secondary minima and overall landscape topologies exhibit marked differences. This structural divergence is reminiscent of the unique feature of the A2V mutation, which increases the risk of AD in the homozygous state but remains non-pathogenic in heterozygous carriers, may reflect the reduced conformational compatibility between WT and A2V variants. For both A2T and WT_1−6D_ variants, their free energy landscapes and the specific locations of minima diverge significantly from the WT. (Figure 3(c,d)) This structural departure is consistent with experimental observations that characterize A2T and WT_1−6D_ as having a protective role against AD. Although the shape of A2V_1−6D_ exhibits a morphology similar to that of WT, the locations of its dominant minima are shifted to the right. (Figure 3(e)) For instance, while the dominant Cluster 0 of the WT is centered at (−1, −1), the corresponding Cluster 1 of A2V_1−6D_ is repositioned near (0,-2). In the A2T_C*β*_ variant, the inversion of the second chiral center (C*β*) of Threonine fundamentally shifts its principal minima. Notably, the dominant Cluster 2 is positioned adjacent to the Cluster 0 of WT. (Figure 3 (f)) Comparing the landscape with the A2T one, we hypothesized that the protective role of the A2T mutation may be diminished with such a stereochemical alteration.

To further quantify the similarity between these clusters, a writhe analysis ^69^ capable of capturing the crossings of curves in 3D space, was employed here. The only parameter that can be tuned is the segment length (*l*) which describes the offset of adjacent C*α* atoms (C*α*_*i*_-C*α*_*i*+*l*_). A range of segment lengths (*l* ∈ {1,…, 5}) was evaluated across the various conformational clusters of the A*β* variants. Literature has established that the discrete polygonal chain representing the protein backbone can be smoothed significantly by increasing the segment length (*l*). ^70^ Specifically, shorter segment lengths are ideal for capturing fine-grained local structures, whereas longer segments excel at resolving global rearrangements and large-scale conformational changes. To this end, multiscale writhe features as suggested ^69^ were used here. Given that the writhe values exhibited a consistent trend as *l* increased from 2 to 5, we only include segment length of 1 and 5 for the similarity estimation. Comprehensive results detailing writhe features of different segment lengths are provided in the Supporting Information (Figures S14–S19). The metric for defining the similarity between any two clusters is provided in Eq. S4.

As detailed in Figure 3(g), Cluster 0 of the WT exhibits the highest similarity to Cluster 0 of the A2V variant, which is consistent with the topographical overlap observed in their respective free energy landscapes (Figure 3 (a,b)). Although high similarity was also observed between the Cluster 0 of A2T and Cluster 0 of WT, it is primarily attributed to the broader region of Cluster 0 identified by the clustering algorithm (Figure S13(b)). In contrast, the similarity scores for WT_1−6D_ and A2V_1−6D_ are significantly lower than those observed for A2V. This is particularly evident for WT_1−6D_, reflecting a more pronounced structural divergence from the wild type ensemble. When closely examining the similarity by A2T_C*β*_ and A2T, we can see that similarity of A2T_C*β*_ increases a lot compared to ones of A2T, particularly for the Cluster 2 of A2T_C*β*_. All these data further confirm the reliability of the free energy landscape we constructed and demonstrate the ability of writhe features to discriminate conformational state of IDPs. A detailed similarity value can be found in Table S3.

### 3.4 Mutation and Chirality Induced Structural Changes

The detailed structural conformations are characterized and summarized in Figure 4. From these results, it is evident that Cluster 0 of the WT and Cluster 0 of the A2V variant exhibit striking similarities in their curve crossings, consistent with the similarity score provided in Figure 3 (g) and Table S3. Beyond static structural comparisons, a closer examination of the secondary structure populations derived from the conformations within the two clusters further underscores their similarity. As shown in Figures S20 and S21, both clusters exhibit pronounced *β*-strand populations at the N-and C-terminal regions, accompanied by *α*-helical propensity in the central residues. In both cases, the N-and C-termini are sequestered beneath the CHC (a transition region colored green to yellow), which consequently leaves the hydrophobic region itself exposed to the solvent and may increase the thermodynamic driving force for intermolecular association and aggregation. In addition, the CHC region is exposed across all three clusters identified in the WT. However, in Cluster 1 of the A2V variant, which comprises 5 % of the ensemble, the CHC region is clamped between N-and C-termini. The secondary structure population of Cluster 1 of A2V reveals a marked increase in strand content within the CHC region; this transition can be attributed to the tight spatial constraints imposed by the flanking termini, which effectively stabilize the strand conformation through increased local packing. (Figure S21) Despite its low occupancy, this sub-population in accordance with Ostwald’s rule of stages ^71^ may represent a key metastable intermediate and potentially explaining the differential aggregation behaviors observed experimentally in homozygous and heterozygous A2V carriers.

**Fig. 4.**
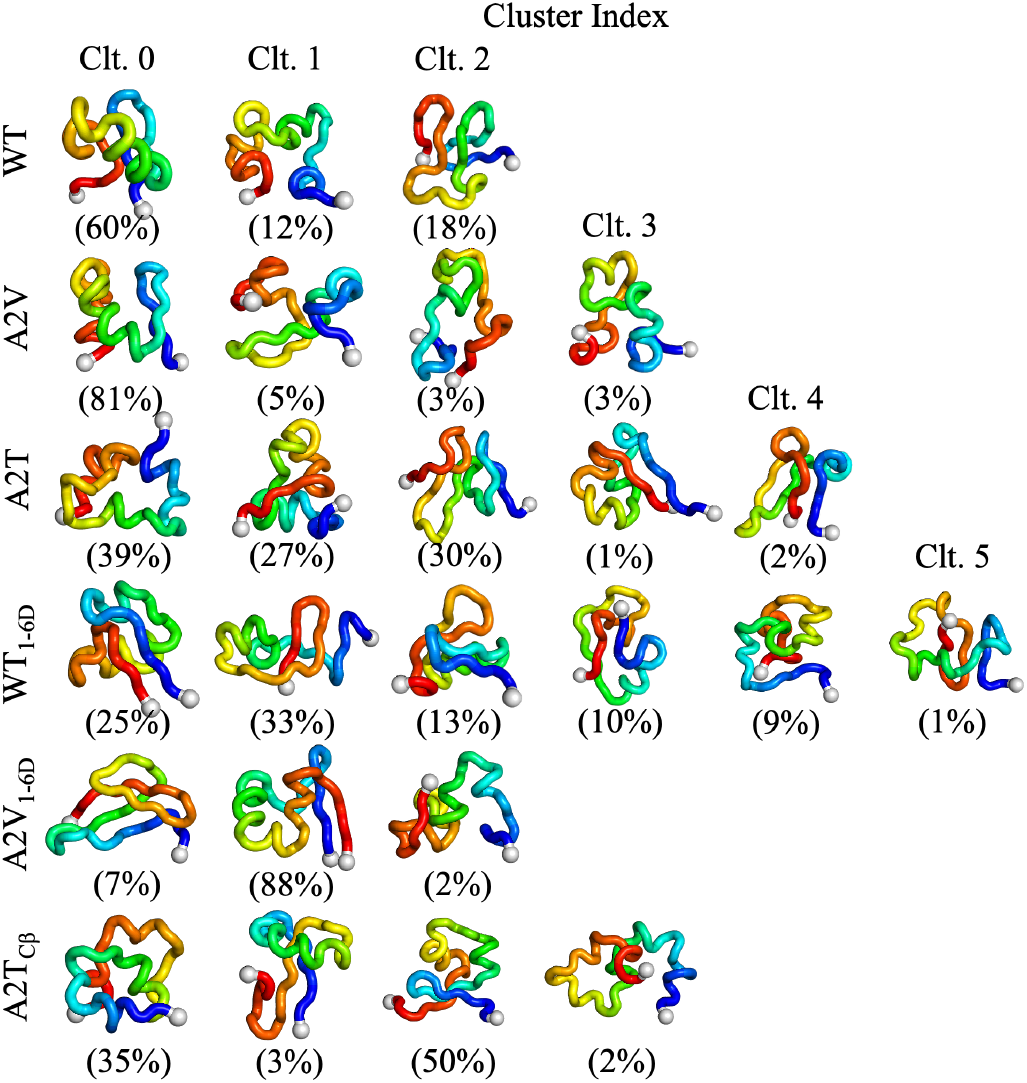
Representative structure of clusters identified across six A*β* variants. All structures are rendered in the ribbon style, with the backbone colored in a transition from blue (N-terminus) to red (C-terminus). The C*α* atoms of terminal residues are indicated by gray sphere. The population percentages of each cluster, as determined by DBSCAN clustering, are provided in parentheses.

Furthermore, in variants exhibiting protective effects, specifically A2T, WT_1−6D_, and A2V_1−6D_, we identified conformations characterized by parallel interactions between the N-and C-termini. This specific topological arrangement brings the termini into close proximity, resulting in a more compact and folded conformation. The population of these compact conformations varies across the protective variants. They constitute 3 % of the total clusters in A2T (comprised of Clusters 3 and 4), 25 % of the population in WT_1−6D_ (Cluster 0), and a significant 88 % of the population in A2V_1−6D_ (Cluster 1). Excluding Cluster 1 of A2V_1−6D_, these clusters possess extremely high strand population in their N- and C-terminal regions. (Figure S22 - S24) This structural enrichment is driven by the tight intramolecular coupling between the two termini. In addition to the parallel arrangement, WT_1−6D_ exhibits an anti-parallel mode in Clusters 1, 3, 4, and 5, accounting for 33 %, 10 %, 9 %, and 1 % of the population, respectively. Such a structural feature is also detected in A2T_C*β*_, although with only 3 % population in Cluster 1. This phenomenon can be attributed to the alteration of chirality, which restricts the conformational space accessible to the backbone, thereby enhancing rigidity and favoring the formation of compact, self-shielded structures. The secondary structure profile of A2T_C*β*_ (Cluster 2) resembles the WT (Cluster 0) more closely than the original A2T variant, highlighting a loss of protective structural traits and a potential shift toward a pathogenic state induced by the C*β* chirality change. (Figure S25)

### 3.5 Phase Transition Derived from Temperature Mapping

All preceding analyses focused on a specific temperature, however, additional insights can be deduced from temperature gradients, such as the entropy ^72^ and boundary of phase transitions ^73^. To complement the structural analysis with thermodynamic insights, phase transitions deduced from temperature-dependent simulations were presented below. Here two properties, namely, the radius of gyration (*R*_*g*_) and heat capacity (*C*_*v*_) were analyzed. Details of the heat capacity calculations were provided in Eq. S3.

As shown in Figure 5(a), the plot of *δR*_*g*_*/δT* as a function of *T* for the WT peptide exhibits a sharp peak around 355 *K*, indicating the temperature of maximum structural instability. In the pathogenic A2V variant, which increases the risk of Alzheimer’s disease in a homozygous condition, its peak shifts to a higher temperature of approximately 370 *K*. Conversely, for the protective A2T variant, the peak shifts to a lower temperature of roughly 340 *K*. These results indicate A2V is more resistant to temperature effects, whereas A2T is more sensitive. When examining the corresponding chirality modification, a bump appears at around 320 *K* in the WT_1−6D_ variant. Similarly, the peak for A2V_1−6D_ shifts toward a lower temperature compared to A2V. Together, these results revealed that chirality modification at the first six residues renders A*β* more sensitive to temperature, likely due to alterations in backbone rigidity. Interestingly, altering the second chiral center (C*β*) of the Threonine residue (A*β*_*Cβ*_) causes a right-shift in the temperature resistance profile, indicating that this modification imparts a promotive effect compared to the A2T variant. This observation is consistent with our predictions based on the structural and free energy landscape analysis.

**Fig. 5.**
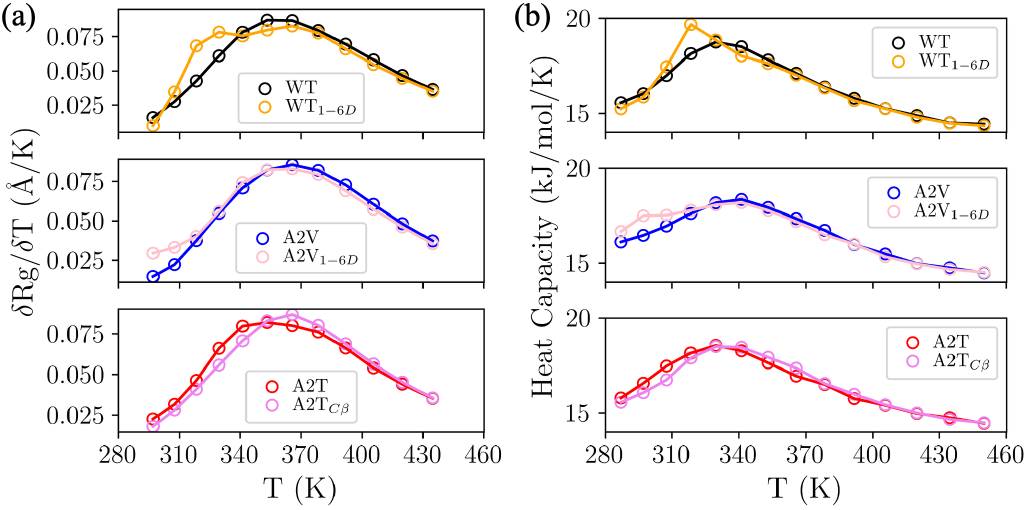
Thermodynamic behavior obtained from replica exchange molecular dynamics. (a) The evolution of the temperature derivative of the radius of gyration (*δR*_*g*_*/δT*) as a function of temperature; (b) Heat capacity (*C*_*v*_) as a function of temperature.

Although the profile of *δR*_*g*_*/δT*(*T*) provides valuable structural insight, the temperature dependence of the heat capacity (*C*_*v*_(T)) provides a more direct measure of the phase transitions ^74–76^ derived from simulations. As shown in Figure 5(b), the general trends derived from *C*_*v*_(T) coincide well with those extracted from *δR*_*g*_*/δT* profile. The peak in the *C*_*v*_(T) profile indicates that the WT system undergoes a significant phase transition at approximately 330 *K*. Below this critical temperature, the peptide predominantly populates a low-energy, collapsed ensemble. In the context of A*β*42, this stable, compact state is widely regarded as the aggregation-prone precursor, whereas temperatures above 330 *K* favor a more entropically driven, disordered state. This temperature is somewhat higher than the one reported by others ^73^. Furthermore, in contrast to the two peaks observed in the heat capacity profile of dimers or oligomers, only a single peak was found in our simulations. This difference is probably attributed to the fact that we simulated the A*β* peptide monomer rather than the oligomeric states reported in literature ^73^. It should also be kept in mind that due to inherent limitations in the implicit solvent model, the simulated transition temperatures do not align precisely with experimental observations. Nevertheless, the calculated trends remain physically meaningful and representative of the protein’s behavior.

## 4 Conclusions

In this work, we investigated the effects of perturbation within the first six N-terminal residues of the A*β*42 peptide, namely, WT, A2V, A2T, WT_1−6D_, A2V_1−6D_, and A2T_*Cβ*_. This included both sequence mutations and chirality modulations at the *Cα* position, as well as the second chiral center (C*β*) of Threonine–a modification that is frequently overlooked in such contexts. Given the intrinsically rugged conformational landscape of the A*β* peptide, temperature-based replica exchange molecular dynamics (T-REMD) simulations were employed to fully explore its diverse conformational ensembles.

First, through analyzing the effect size (*E*) and population change (|Δ%|) of each residue within *ϕ* /*ψ* space, we observed that the effects of mutation and chirality modulation are not solely localized at the N-terminus where they are incorporated. Instead, these perturbations exhibit a specific long-distance effect. Notably, the regions most affected by such modifications fall into the CHC region, a domain previously demonstrated to be responsible for fibril maturation and a primary target for clinical trial interventions. Furthermore, we examined the free energy landscapes projected onto the first two principal components, alongside representative structures extracted from their respective local basins. We concluded that A2V exhibits the highest similarity to WT but differs in the locations and populations of secondary basins. The similarity between A2V and WT may explain the dual effect of A2V that will be protective in the heterozygous state but pathogenic in the homozygous state. For variants that decrease the risk of AD, the topology and location of the energy basins differ significantly from WT, particularly in the case of WT_1−6D_. We further quantified the structural similarities among representative conformations using writhe analysis, which demonstrated that A2V maintains a higher structural similarity to the WT than any other variant. Interestingly, a nuanced alteration in the second chiral center of Threonine (A2T_*Cβ*_) restores structural similarity to the WT, reversing the trend observed in the A2T variant.

Beyond analyzing thermodynamic properties at specific temperatures, we also investigated behaviors across thermal gradients, which are more directly related to the phase transitions of the system. Taking two metrics into consideration, namely, the temperature derivative of the radius of gyration (*δR*_*g*_*/δT*) and heat capacity (*C*_*v*_(T)), the transition peaks shift toward higher temperatures for protective variants and lower temperatures for the pathogenic variant were observed. This upward shift in transition temperature indicates that the system is more thermally resilient, maintaining its structural integrity at higher temperatures. We also examined the impact of A2T_C*β*_ modification on aggregation propensity. Consistent with our free energy landscape analysis, both the peaks of *δR*_*g*_*/δT* and *C*_*v*_(T) exhibit a rightward shift toward higher temperatures, indicating their increased aggregation propensity.

These findings suggest that the A*β* N-terminus is a viable site for therapeutic intervention and highlight the use of chiral inversion as an innovative protocol for modulating the structural stability and aggregation pathways of pathogenic proteins.

## Author contributions

Qiang Zhu: conceptualization, validation, visualization, investigation, formal analysis, and writing – original draft preparation, funding acquisition; Haibo Yu: conceptualization, supervision, projection administration, funding acquisition, and writing – reviewing and editing.

## Data availability

All data and scripts used in this work are available at https://github.com/QZResearch/Abeta42.

## Acknowledgements

We thank Dr Qinghua Liao (University of Barcelona, Spain) for insightful comments on the draft. We thank supports provided by the Australian Research Council Centre of Excellence in Quantum Biotechnology through project number CE230100021. Q.Z. is supported by UOW Vice-Chancellor’s Research Fellowship. This project was undertaken with the assistance of resources and services from the National Computational Infrastructure (NCI), which is supported by the Australian Government through the National Computational Merit Allocation Scheme (NCMAS project eh83 & v15). This work was also supported by resources provided by the Pawsey Supercomputing Research Centre’s Setonix Super-computer with funding from the Australian Government and the Government of Western Australia.

