## supporting information for "N-terminal Chirality and Sequence Variations Modulate the Conformational Landscape of Amyloid-beta 42^†^"

### Contents

**S1 Summary of the simulated systems** **S4**

**S2 Convergence test and performance estimation** **S5**

S2.1 Evolution of secondary structure as a function of simulation time . . . . . . S5

S2.2 Radius of gyration ($R_g$) . . . . . . . . . . . . . . . . . . . . . . . . . . . . S6

S2.3 Evolution of secondary structure as a function of temperature . . . . . . . . S7

S2.4 Chemical shifts . . . . . . . . . . . . . . . . . . . . . . . . . . . . . . . . . S8

**S3 Ramachandran plots** **S9**

S3.1 WT . . . . . . . . . . . . . . . . . . . . . . . . . . . . . . . . . . . . . . . S9

S3.2 A2V . . . . . . . . . . . . . . . . . . . . . . . . . . . . . . . . . . . . . . . S10

S3.3 A2T . . . . . . . . . . . . . . . . . . . . . . . . . . . . . . . . . . . . . . . S11

S3.4 $A2V_{1-6D}$ . . . . . . . . . . . . . . . . . . . . . . . . . . . . . . . . . . . S12

S3.5 $WT_{1-6D}$ . . . . . . . . . . . . . . . . . . . . . . . . . . . . . . . . . . . S13

S3.6 $A2T_{C\beta}$ . . . . . . . . . . . . . . . . . . . . . . . . . . . . . . . . . . . S14

**S4 Conformational sampling change in first 6 residues** **S15**

**S5 Cluster analysis** **S16**

S5.1 K-distance graph for parameter selection . . . . . . . . . . . . . . . . . . . S16

S5.2 Summary of the chosen parameters for cluster analysis . . . . . . . . . . . S17

S5.3 Clustering results mapped onto PCs 1 and 2 . . . . . . . . . . . . . . . . . S18

S5.4 Strategy of selecting representative structures . . . . . . . . . . . . . . . . S18

**S6 Heat capacity** **S19**

**S7 Writhe analysis** **S20**

S7.1 WT . . . . . . . . . . . . . . . . . . . . . . . . . . . . . . . . . . . . . . . S20

S7.2 A2T . . . . . . . . . . . . . . . . . . . . . . . . . . . . . . . . . . . . . . . S21

S7.3 A2V . . . S22

S7.4 $WT_{1-6D}$ . . . S23

S7.5 $A2T_{C\beta}$ . . . S24

S7.6 $A2V_{1-6D}$ . . . S25

S7.7 Similarity between clusters using the writhe feature as input . . . S26

**S8 Secondary structure analysis of each cluster across 6 variants** **S27**

S8.1 WT . . . S27

S8.2 A2V . . . S28

S8.3 A2T . . . S29

S8.4 $WT_{1-6D}$ . . . S30

S8.5 $A2V_{1-6D}$ . . . S31

S8.6 $A2T_{C\beta}$ . . . S32

**References** **S33**

### S1 Summary of the simulated systems

Table S1: Summary of the variants of systems studied in this work together with their descriptions and accumulated simulation time.

| No. | Name | Description | $N_{replica}$ | Time ($\mu s$) |
|---|---|---|---|---|
| 1 | WT | wild-type amyloid beta 42 (A$\beta$42) | 14 | 11.2 |
| 2 | A2T | A$\beta$42 with the second residue mutated from A (Ala) to T (Thr) | 14 | 16.8 |
| 3 | A2V | A$\beta$42 with the second residue mutated from A (Ala) to V (Val) | 14 | 11.2 |
| 4 | $\mathrm{WT}_{1-6\mathrm{D}}$ | A$\beta$42 with the chirality of the first 6 residues changed from L to D | 14 | 16.8 |
| 5 | $\mathrm{A2T}_{\mathrm{C}\beta}$ | A$\beta$42 with the second residue mutated from A to T and the inverted C$\beta$'s chirality of Thr | 14 | 16.8 |
| 6 | $\mathrm{A2V}_{1-6\mathrm{D}}$ | A$\beta$42 with the second residue mutated from A to V and the inverted chirality of the first 6 residues | 14 | 16.8 |

### S2 Convergence test and performance estimation

#### S2.1 Evolution of secondary structure as a function of simulation time

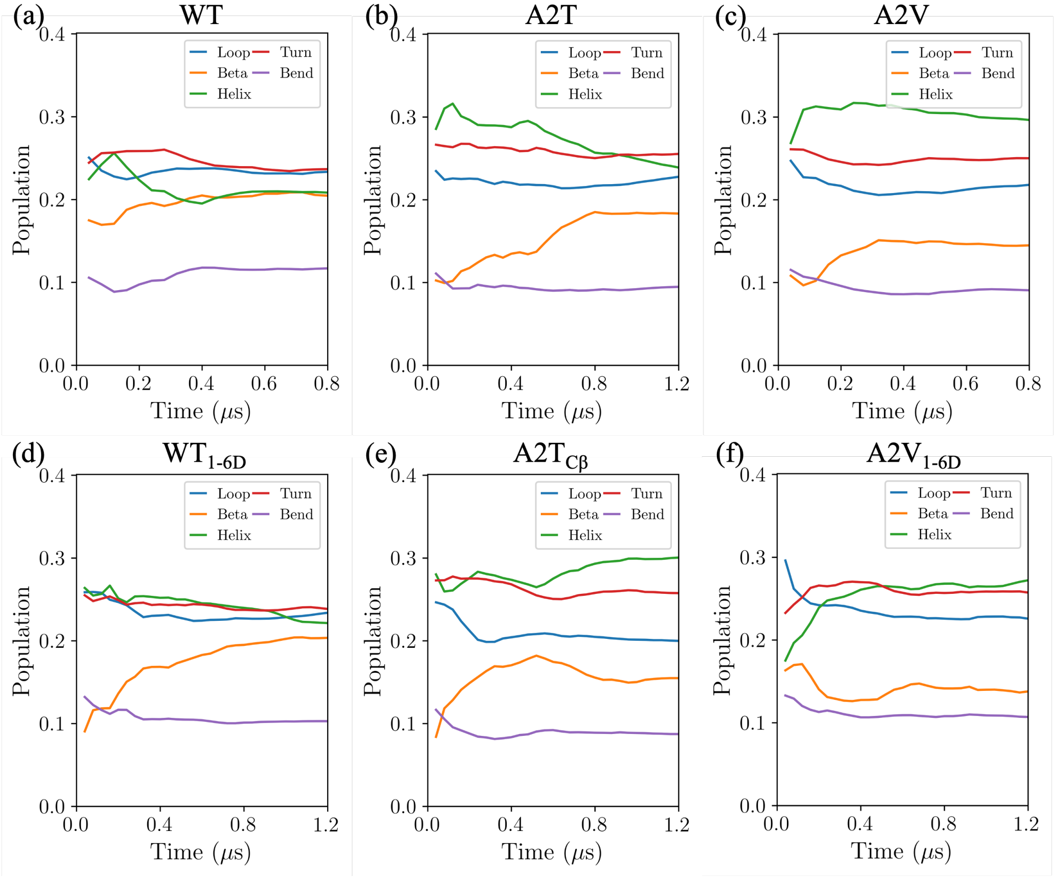


Figure S1: Convergence test of the REMD on the evolution of secondary structure population as a function of simulation time: (a) WT; (b) A2T; (c) A2V; (d) WT$_{1-6D}$; (e) A2T$_{C\beta}$; and (f) A2V$_{1-6D}$, respectively.

#### S2.2 Radius of gyration ($R_g$)

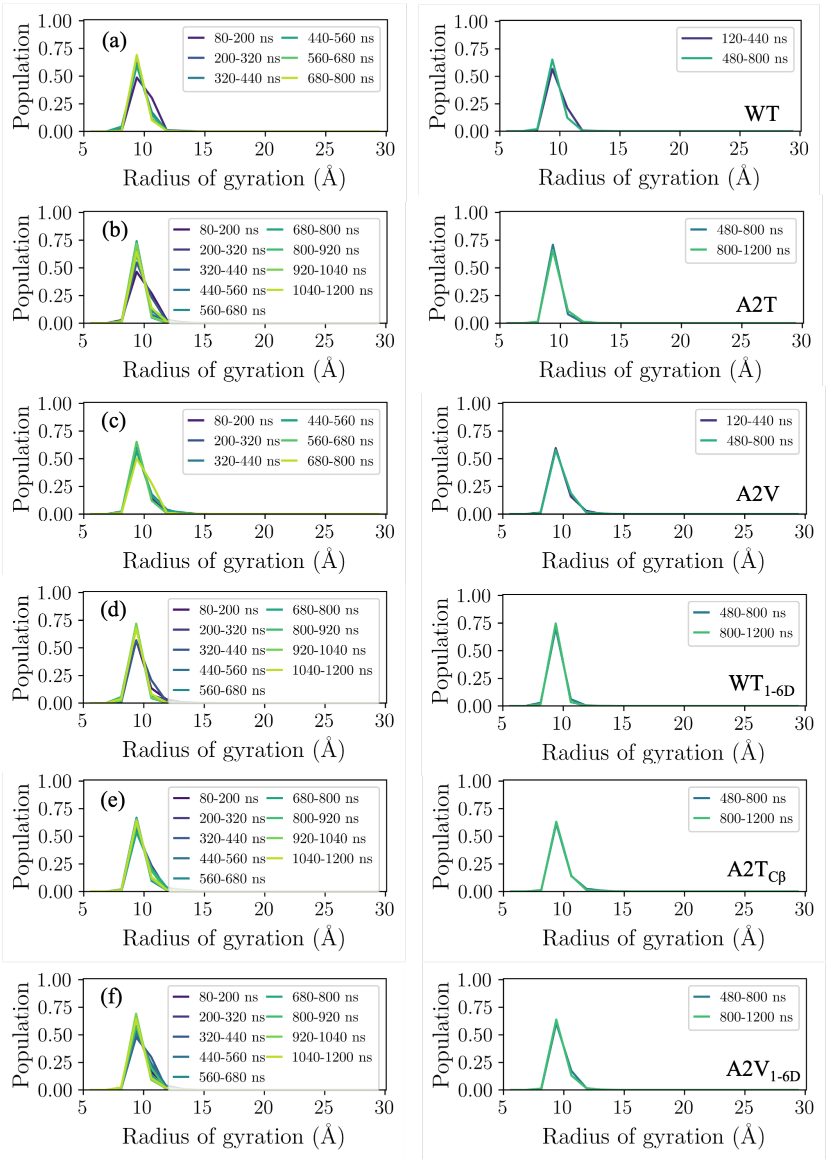


Figure S2: Convergence test of the REMD (T=297.10 $K$ was selected) on the distribution of radius of gyration ($R_g$) of different time intervals, namely, 120 $ns$ (left panel) and 320 $ns$ (400 $ns$ for A2T, A2T$_{C\beta}$, and WT$_{1-6D}$, right panel): (a) WT; (b) A2T; (c) A2V; (d) WT$_{1-6D}$; (e) A2T$_{C\beta}$; and (f) A2V$_{1-6D}$, respectively.

#### S2.3 Evolution of secondary structure as a function of temperature

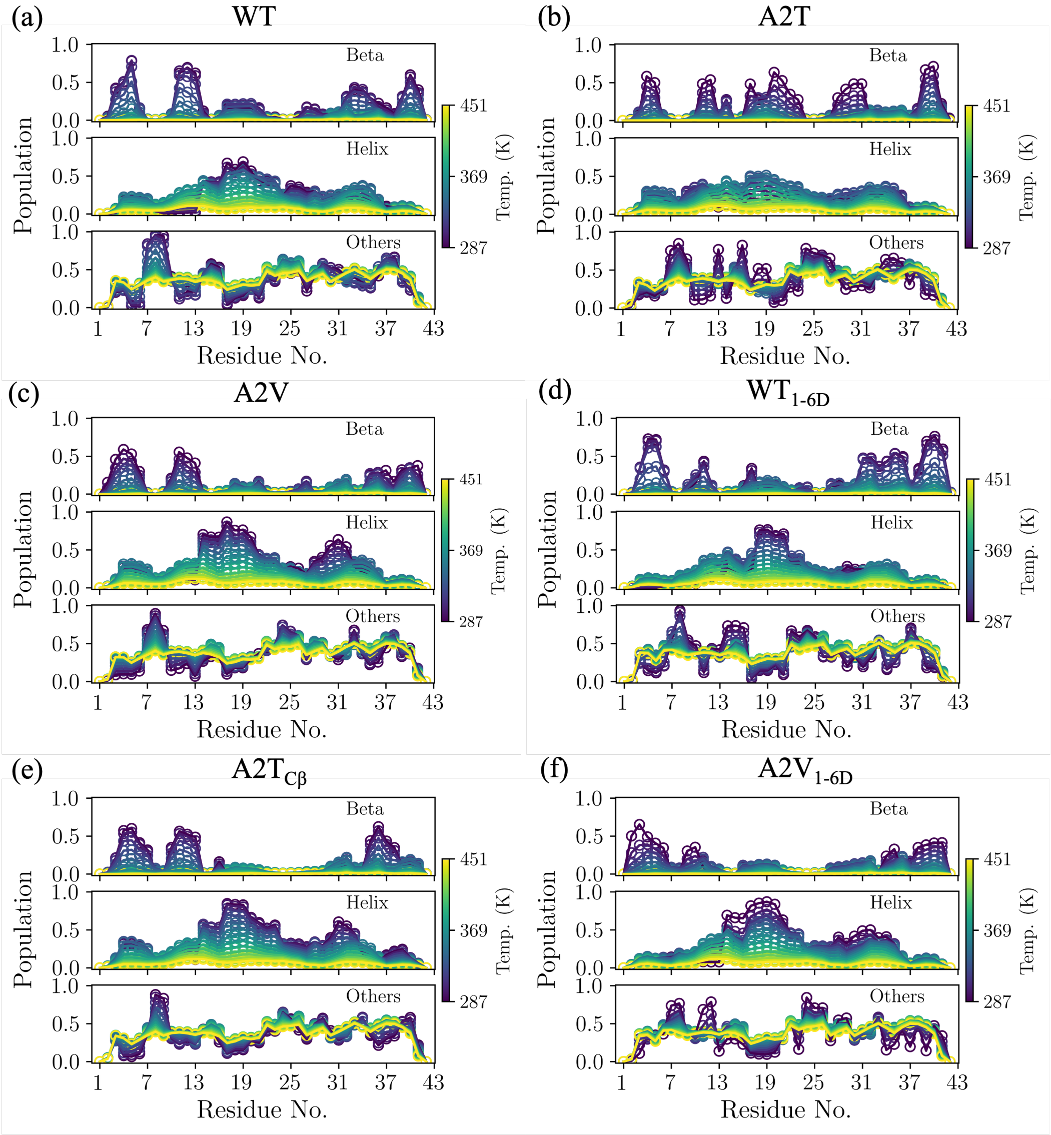


Figure S3: The distribution of secondary structure across 14 temperature replicas: (a) WT; (b) A2T; (c) A2V; (d) $WT_{1-6D}$; (e) $A2T_{C\beta}$; and (f) $A2V_{1-6D}$, respectively. Here beta is composed of extended beta and isolated beta; Helix is composed of 3-10 helix, alpha helix, and Pi(3-14) helix; Others is composed of Turn and Bend.

#### S2.4 Chemical shifts

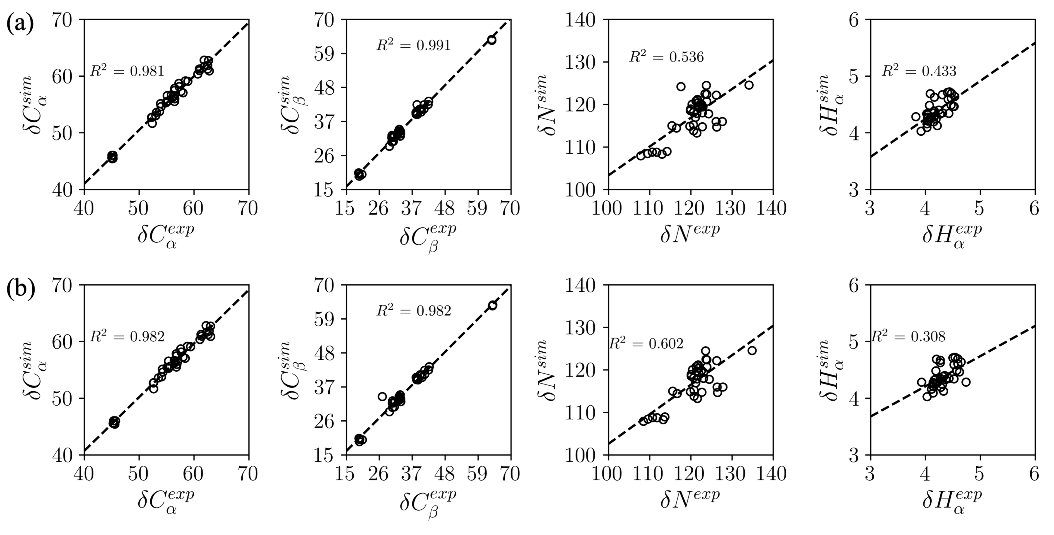


Figure S4: Correlation of simulated and experimental measured chemical shifts. The chemical shift values were calculated using the LEGOLAS[1] methods and conducted on one replica (T=287.0 $K$) of WT. The experimental NMR values were directly extracted from Biological Magnetic Resonance Data Bank (BMRB), corresponding to entries are (a) 25218[2] and (b) 17793,[3] respectively.

### S3 Ramachandran plots

## S3.1 WT

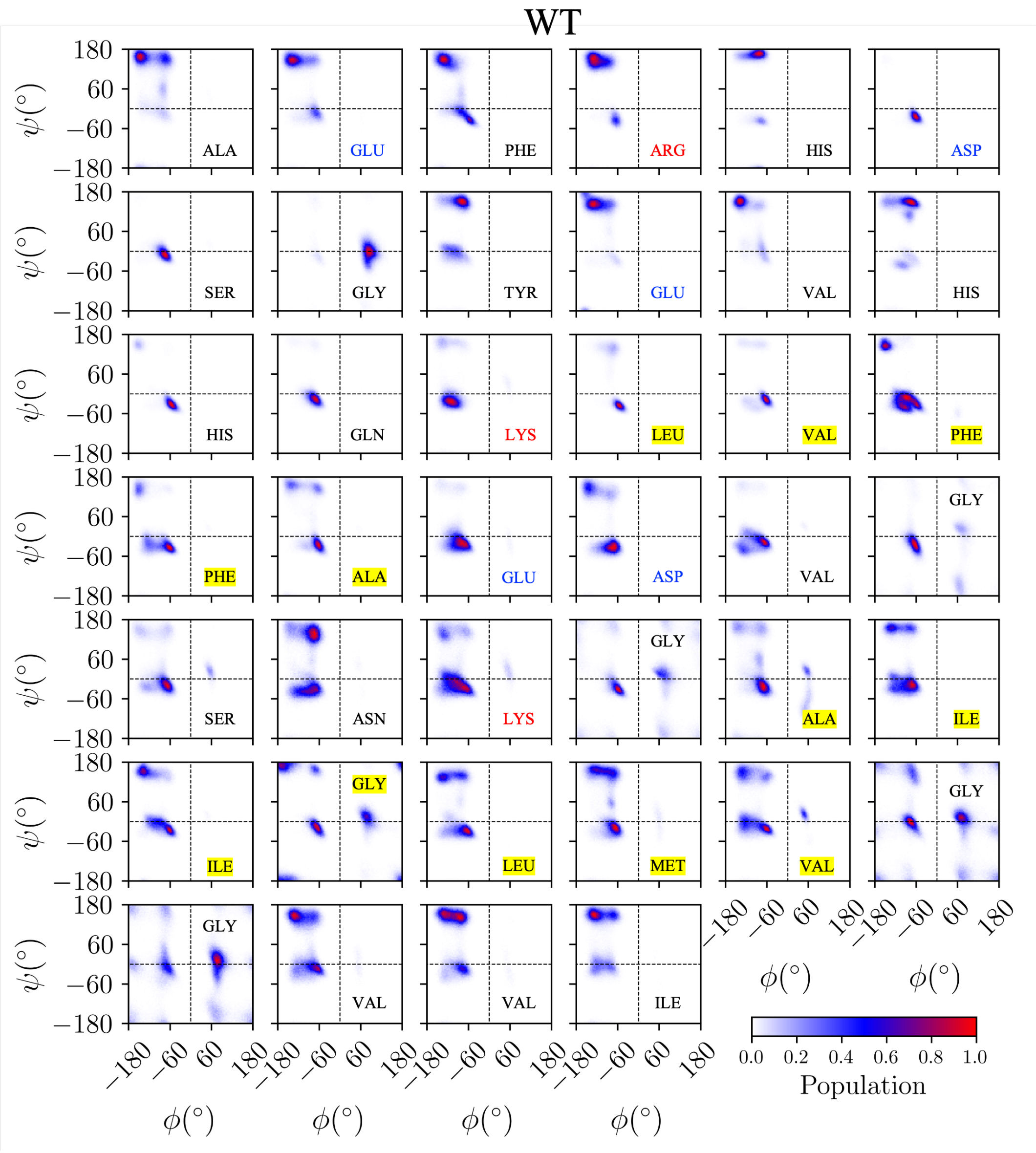


Figure S5: Residue-resolved Ramachandran plots showing the conformational populations sampled by WT.

## S3.2 A2V

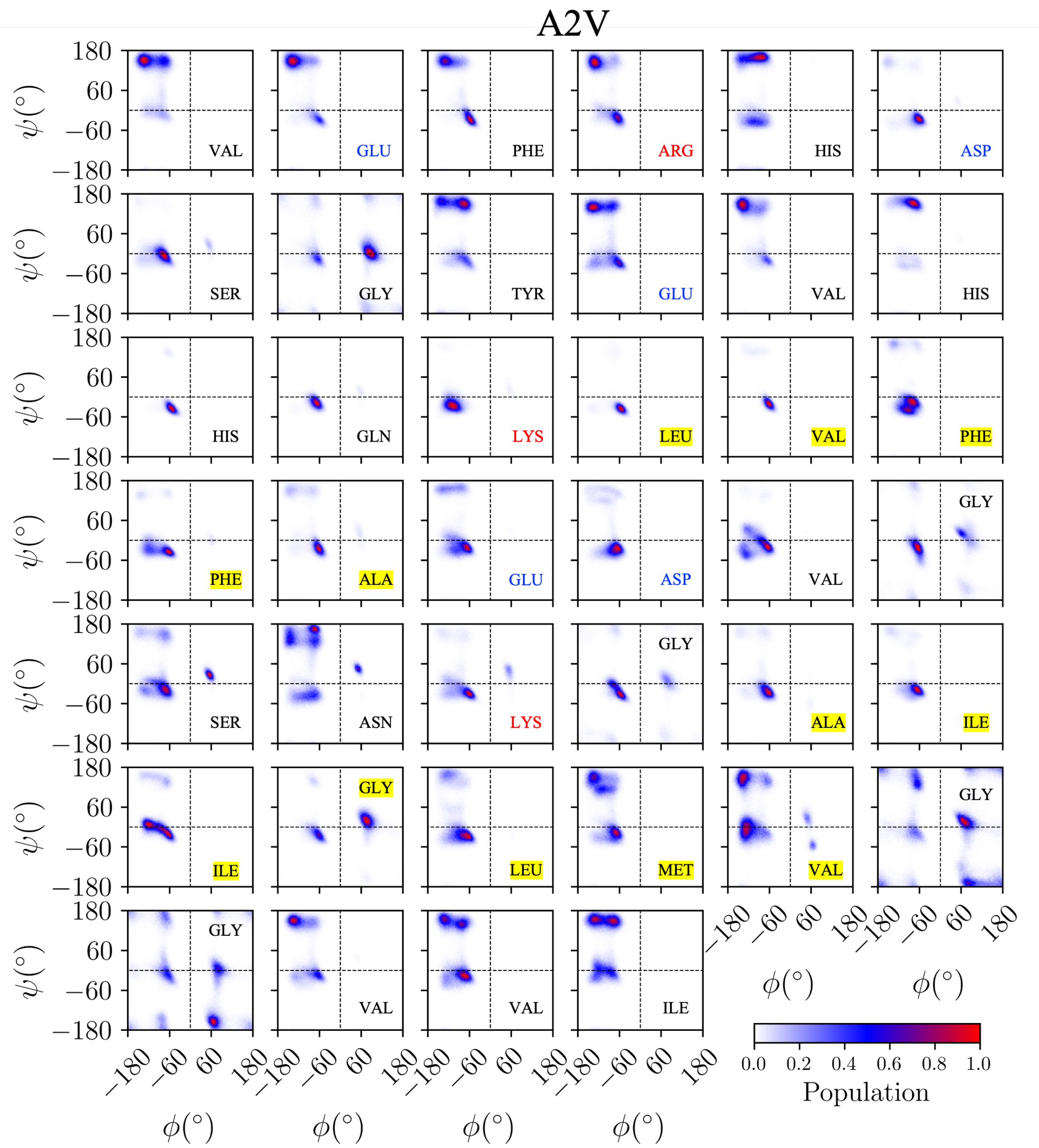


Figure S6: Residue-resolved Ramachandran plots showing the conformational populations sampled by A2V.

## S3.3 A2T

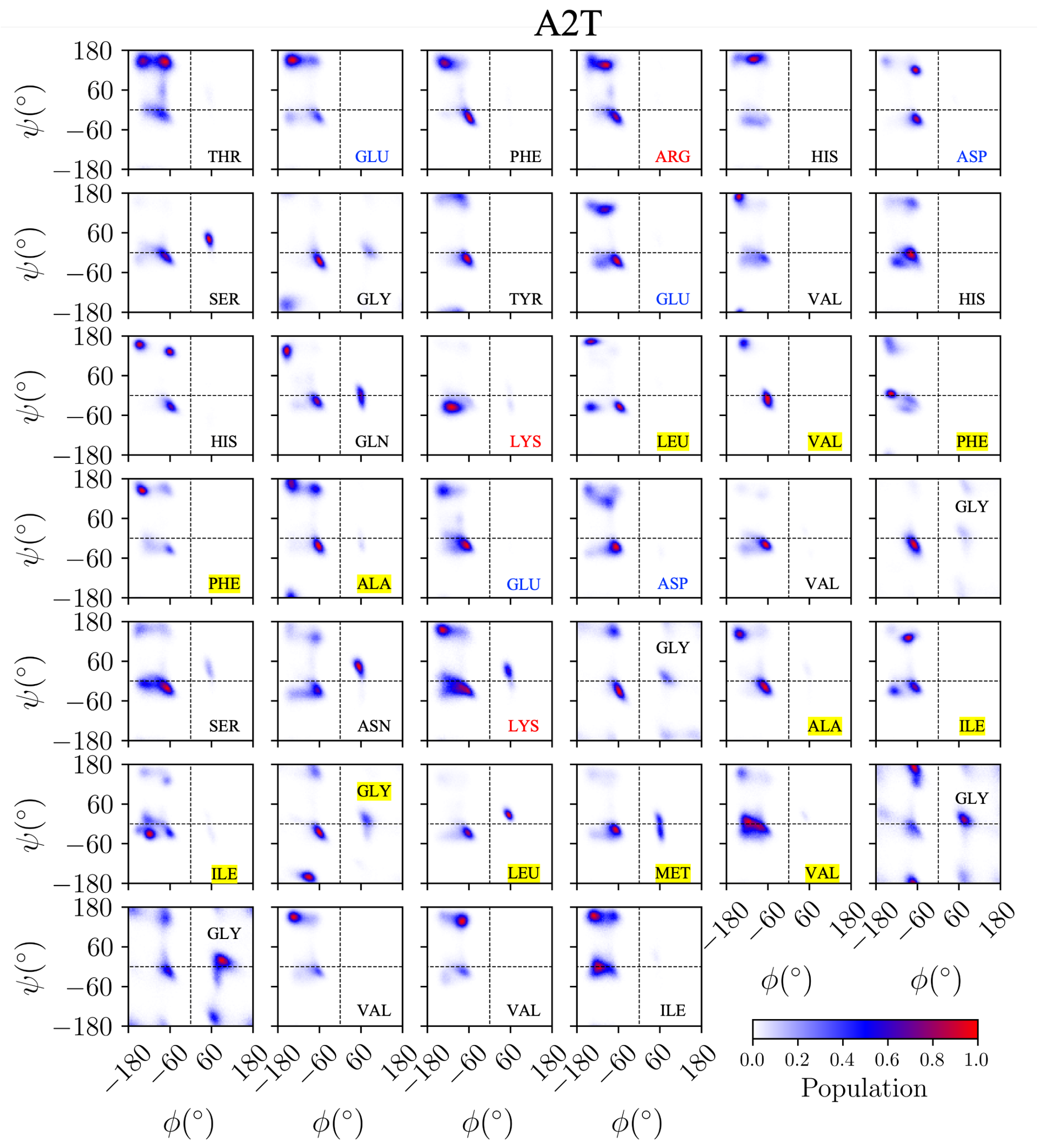


Figure S7: Residue-resolved Ramachandran plots showing the conformational populations sampled by A2T.

## S3.4 $A2V_{1-6D}$

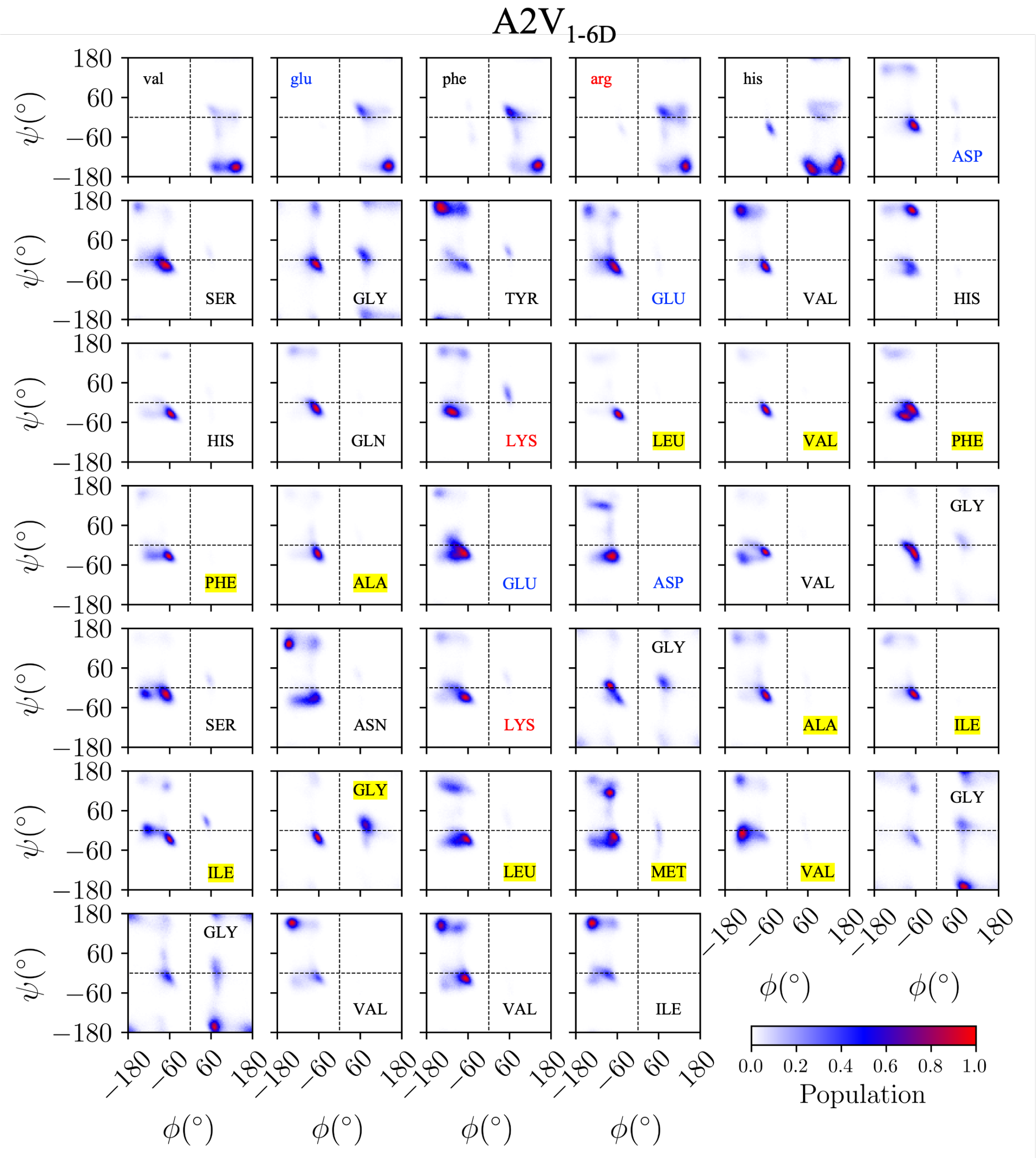


Figure S8: Residue-resolved Ramachandran plots showing the conformational populations sampled by $A2V_{1-6D}$.

##### S3.5 $\mathbf{WT}_{1-6D}$

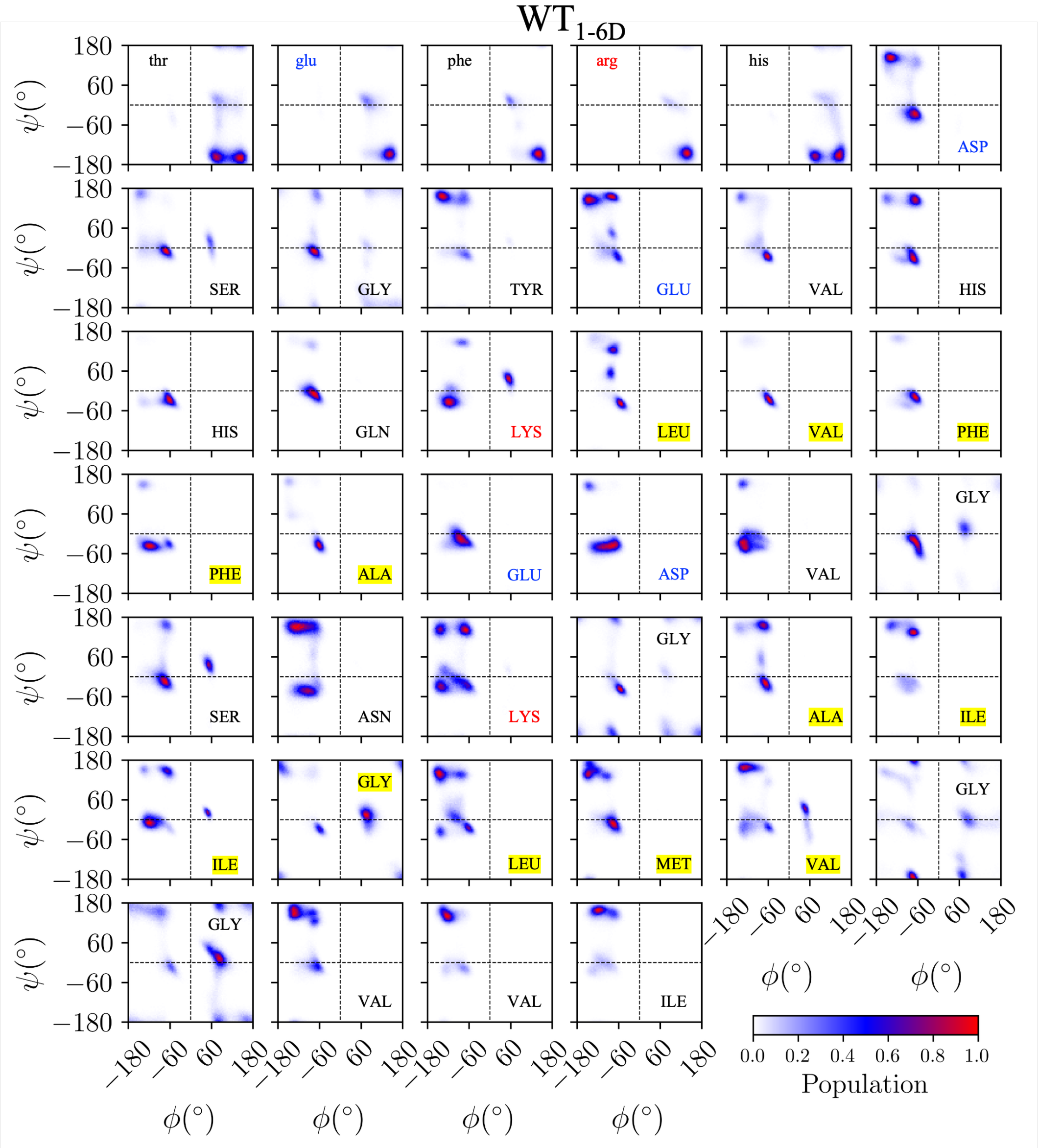


Figure S9: Residue-resolved Ramachandran plots showing the conformational populations sampled by $\mathrm{WT}_{1-6D}$.

#### S3.6 A2T$_{C\beta}$

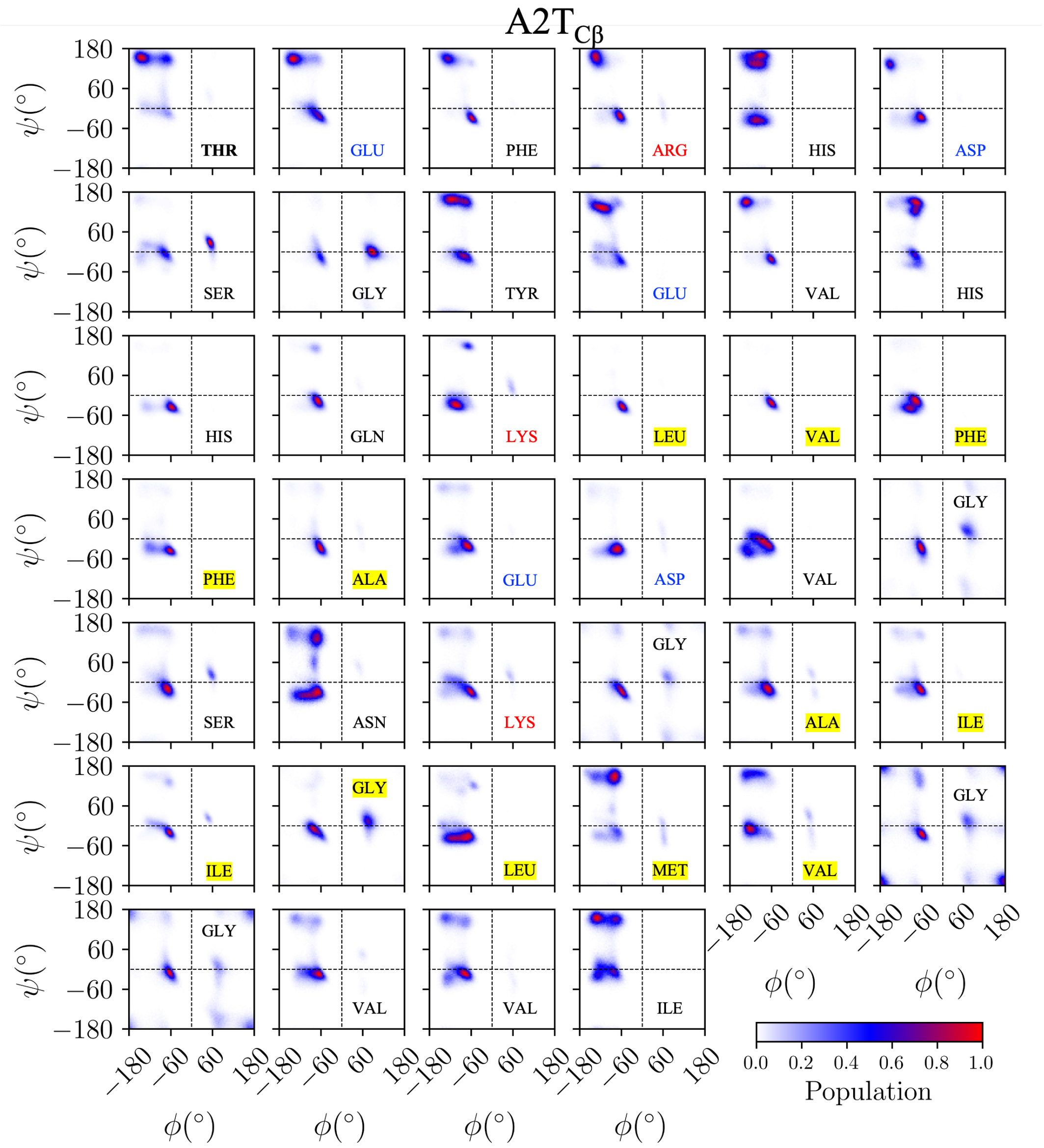


Figure S10: Residue-resolved Ramachandran plots showing the conformational populations sampled by A2T$_{C\beta}$.

#### S4 Conformational sampling change in first 6 residues

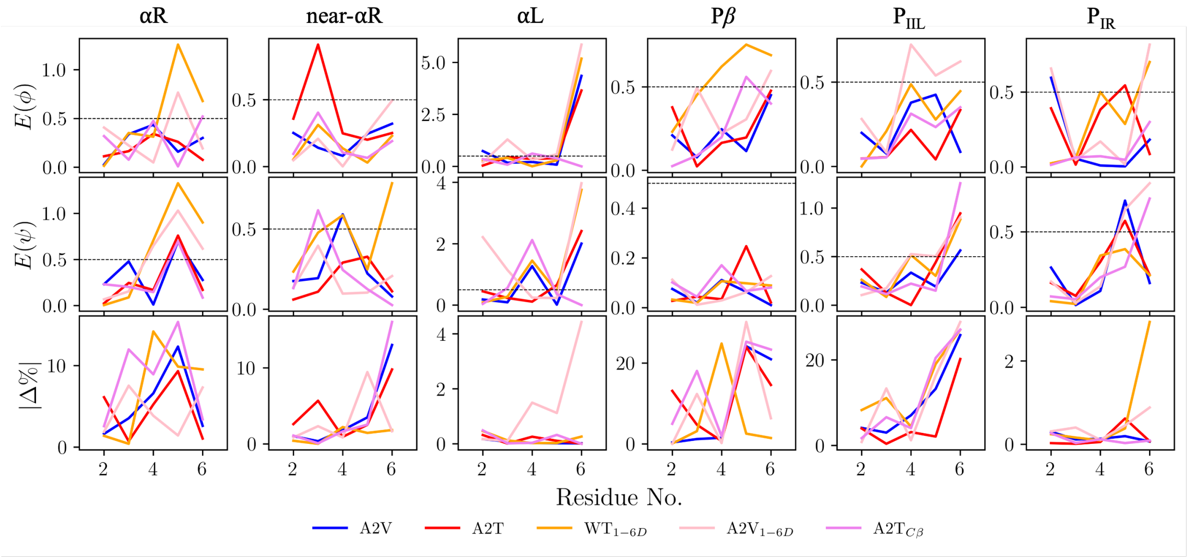


Figure S11: Effect size analysis of the N-terminal hexapeptide. The structural impact within the first six residues is quantified via ($\phi$,$\psi$) dihedral distributions and the absolute percentage change in conformational populations ($|\Delta\%|$) across the investigated A$\beta$ variants with respect to WT. The effect size threshold of 0.5 is indicated by a dashed line.

### S5 Cluster analysis

Here, density-based spatial clustering of applications with noise (DBSCAN) clustering method was employed and implemented with the help of *scikit-learn*.[4] The trajectories were processed with the help of *mdtraj*.[5]

#### S5.1 K-distance graph for parameter selection

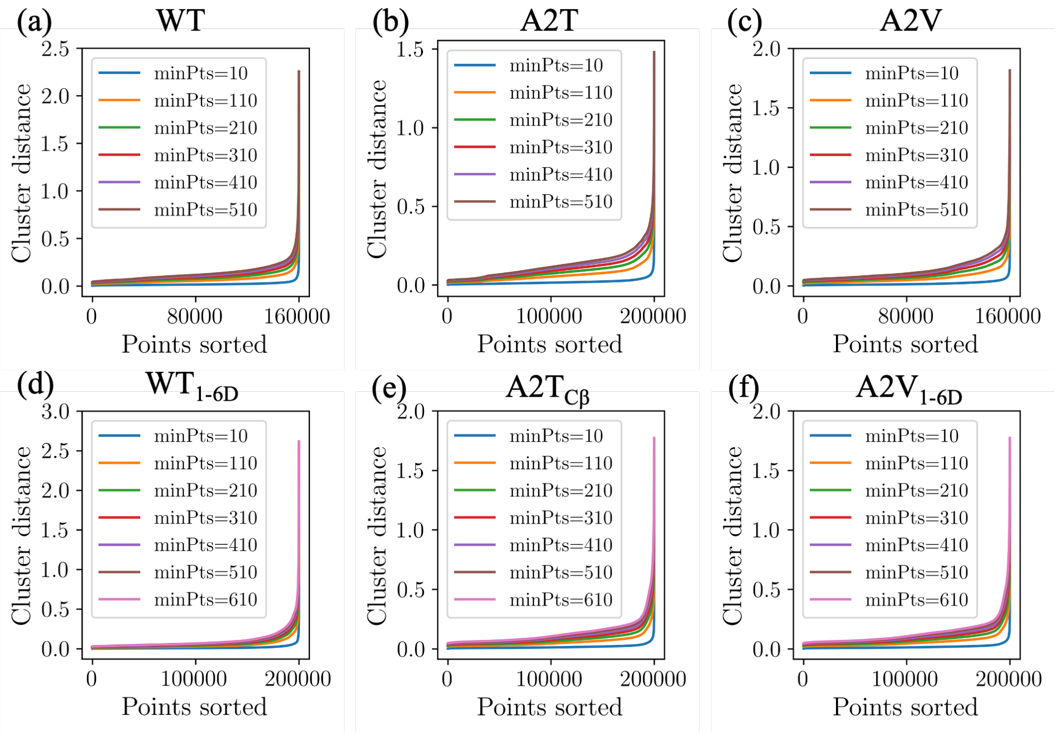


Figure S12: Points sorted by distance to determine the optimized parameter `eps` and `min_samples` for simulations of different variants: (a) WT; (b) A2T; (c) A2V; (d) $\mathrm{WT}_{1-6\mathrm{D}}$; (e) $\mathrm{A2T}_{\mathrm{C}\beta}$; and (f) $\mathrm{A2T}_{1-6\mathrm{D}}$, respectively.

#### S5.2 Summary of the chosen parameters for cluster analysis

Table S2: Summary of two key parameters, namely, eps and min_samples, used in the DBSCAN clustering method.

| Variant | eps | min_samples |
|---|---|---|
| WT | 0.200 | 600 |
| A2T | 0.250 | 400 |
| A2V | 0.200 | 400 |
| $\mathrm{WT}_{1-6\mathrm{D}}$ | 0.100 | 200 |
| $\mathrm{A2T}_{\mathrm{C}\beta}$ | 0.172 | 600 |
| $\mathrm{A2V}_{1-6\mathrm{D}}$ | 0.250 | 600 |

#### S5.3 Clustering results mapped onto PCs 1 and 2

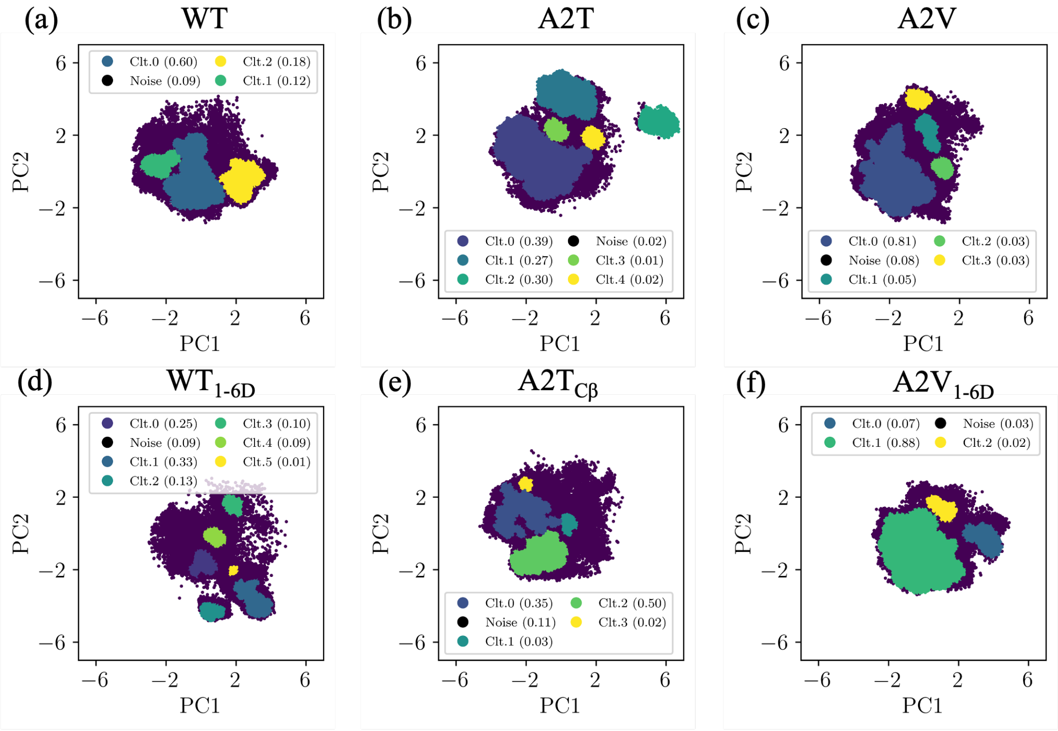


Figure S13: Clusters identified with the help of DBSCAN method and visualized on two principle components (PC1 and PC2). Clusters together with noise were visualized using different colors. Their corresponding populations were indicated in the legend. Detailed parameters were summarized in Table S2.

#### S5.4 Strategy of selecting representative structures

A similarity score $s_{ij}$ was defined according to Eq. S1 to select the centroid as the representative structure within a given cluster.

$$s_{ij} = e^{-d_{ij}/d_{scaled}} \tag{S1}$$

where $d_{ij}$ denotes the pairwise distance between $i_{th}$ and $j_{th}$ points in the transformed PC spaces (top 10 components were retained). The distance is computed using the Euclidean metric. $d_{scaled}$ is the standard deviation of the pairwise distance computed to make the score scale invariant. The representative structure within the individual cluster is picked with the highest similarity and the mathematical expression is defined as below:

$$\arg\max_i \sum_j s_{ij} \tag{S2}$$

### S6 Heat capacity

$$C_v = \frac{\langle E^2 \rangle - \langle E \rangle^2}{k_B T^2} = \frac{\langle \delta E^2 \rangle}{k_B T^2} \tag{S3}$$

where $k_B$ represents the Boltzmann constant; $T$ is the temperature where the system is simulated at. $\langle \cdots \rangle$ is a notation of the ensemble average. $E$ represents the total energy of the system including both potential and kinetic contributions.

### S7 Writhe analysis

## S7.1 WT

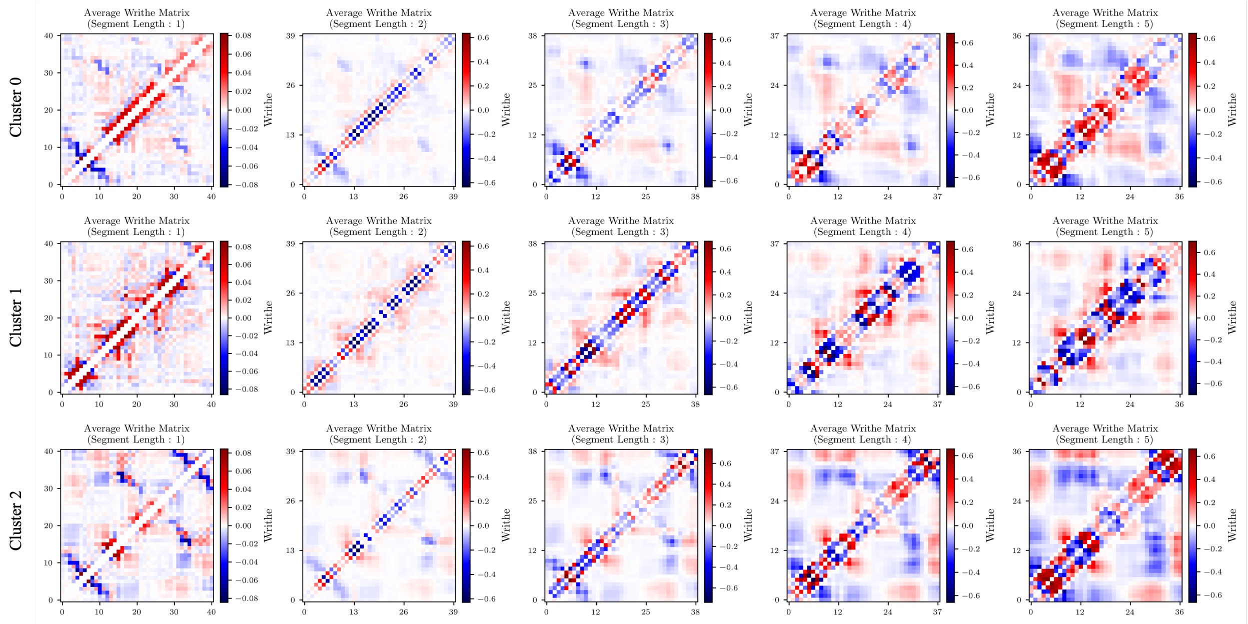


Figure S14: Pairwise writhe matrices computed within selected clusters at different segment length ($l$) ranging from 1 to 5, where the $x$- and $y$-axis indices correspond to protein residue indices of variant, WT.

## S7.2 A2T

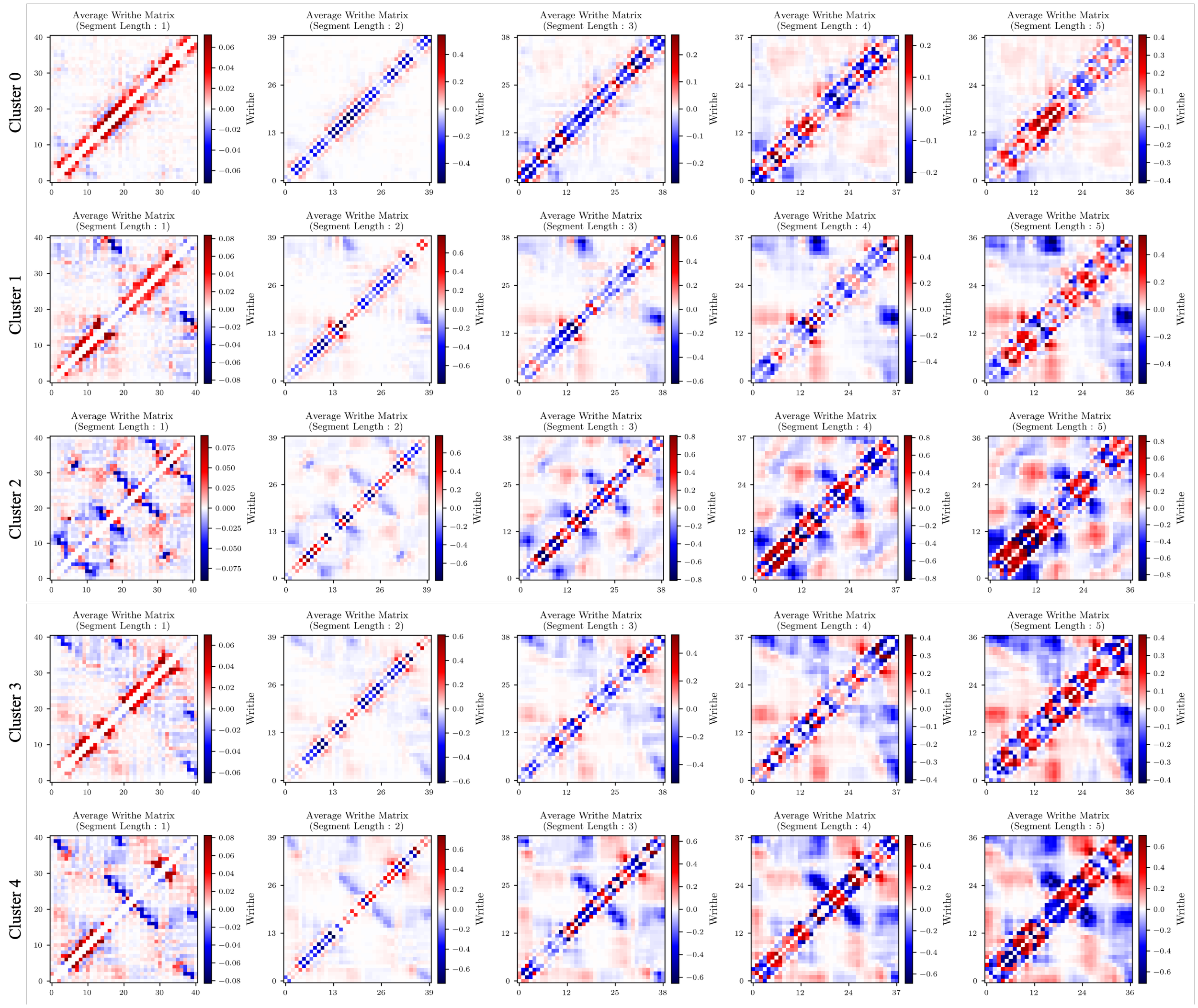


Figure S15: Pairwise writhe matrices computed within selected clusters at different segment length ($l$) ranging from 1 to 5, where the $x$- and $y$-axis indices correspond to protein residue indices of variant, A2T.

## S7.3 A2V

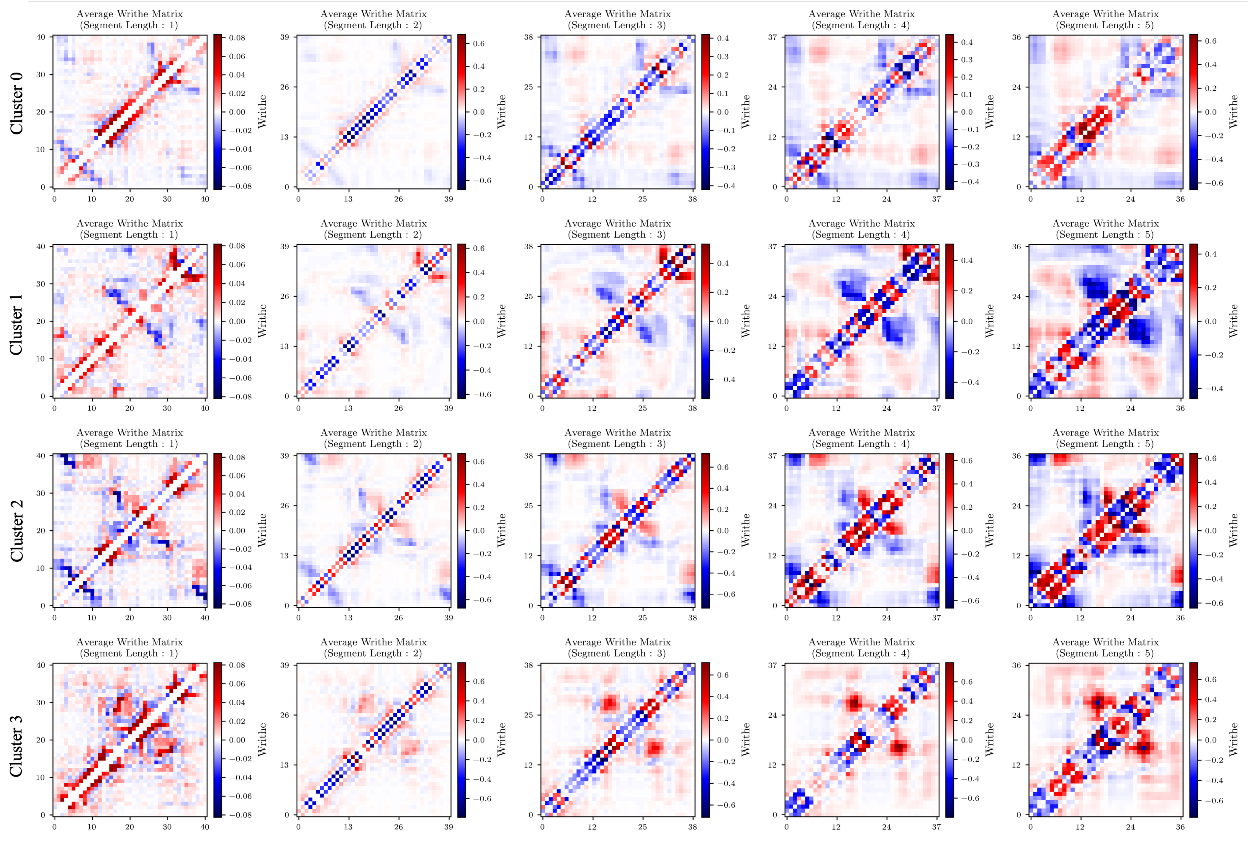


Figure S16: Pairwise writhe matrices computed within selected clusters at different segment length ($l$) ranging from 1 to 5, where the $x$- and $y$-axis indices correspond to protein residue indices of variant, A2V.

#### S7.4 $\mathrm{WT}_{1-6\mathrm{D}}$

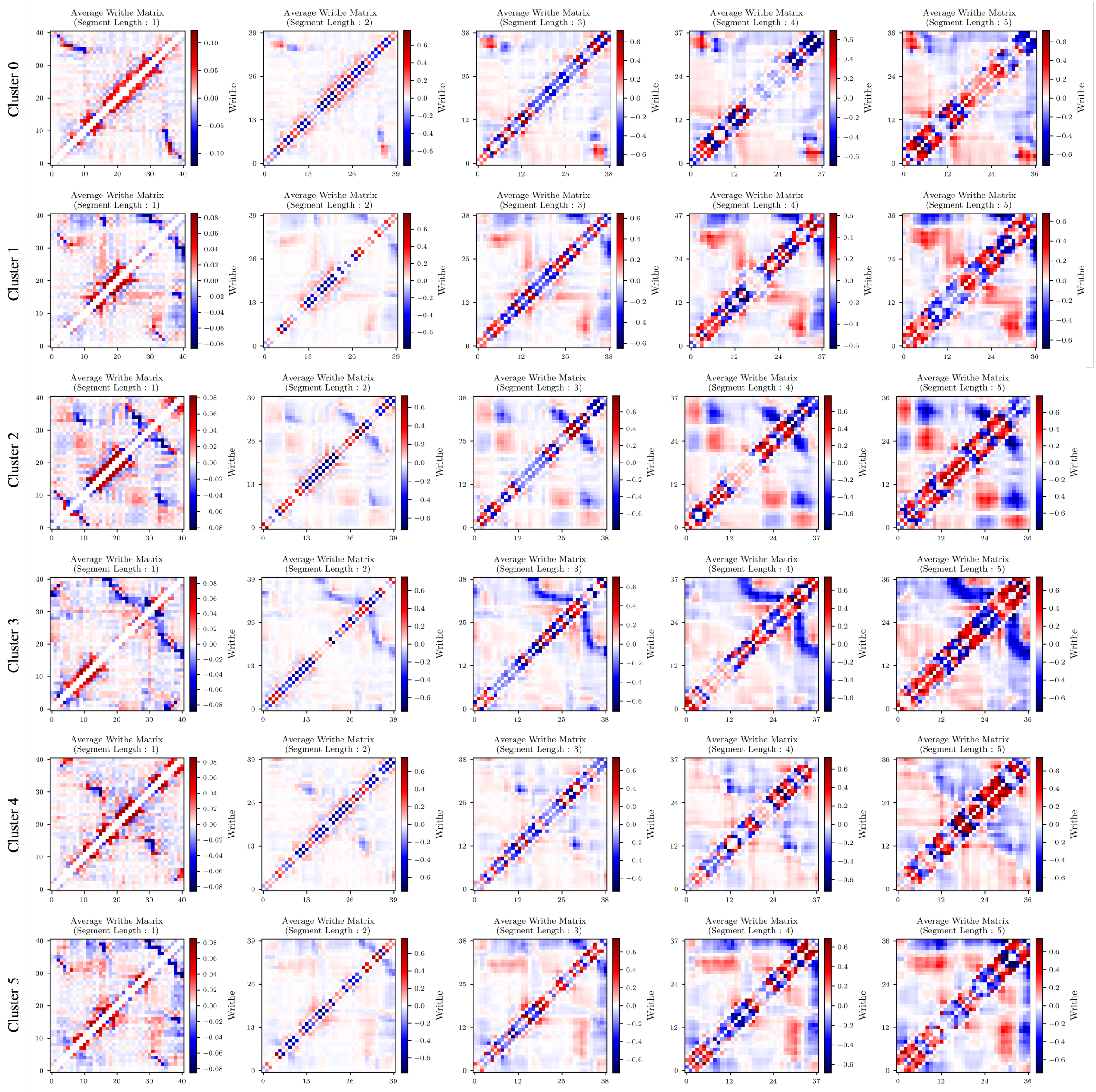


Figure S17: Pairwise writhe matrices computed within selected clusters at different segment length ($l$) ranging from 1 to 5, where the $x$- and $y$-axis indices correspond to protein residue indices of variant, $\mathrm{WT}_{1-6\mathrm{D}}$.

#### S7.5 $A2T_{C\beta}$

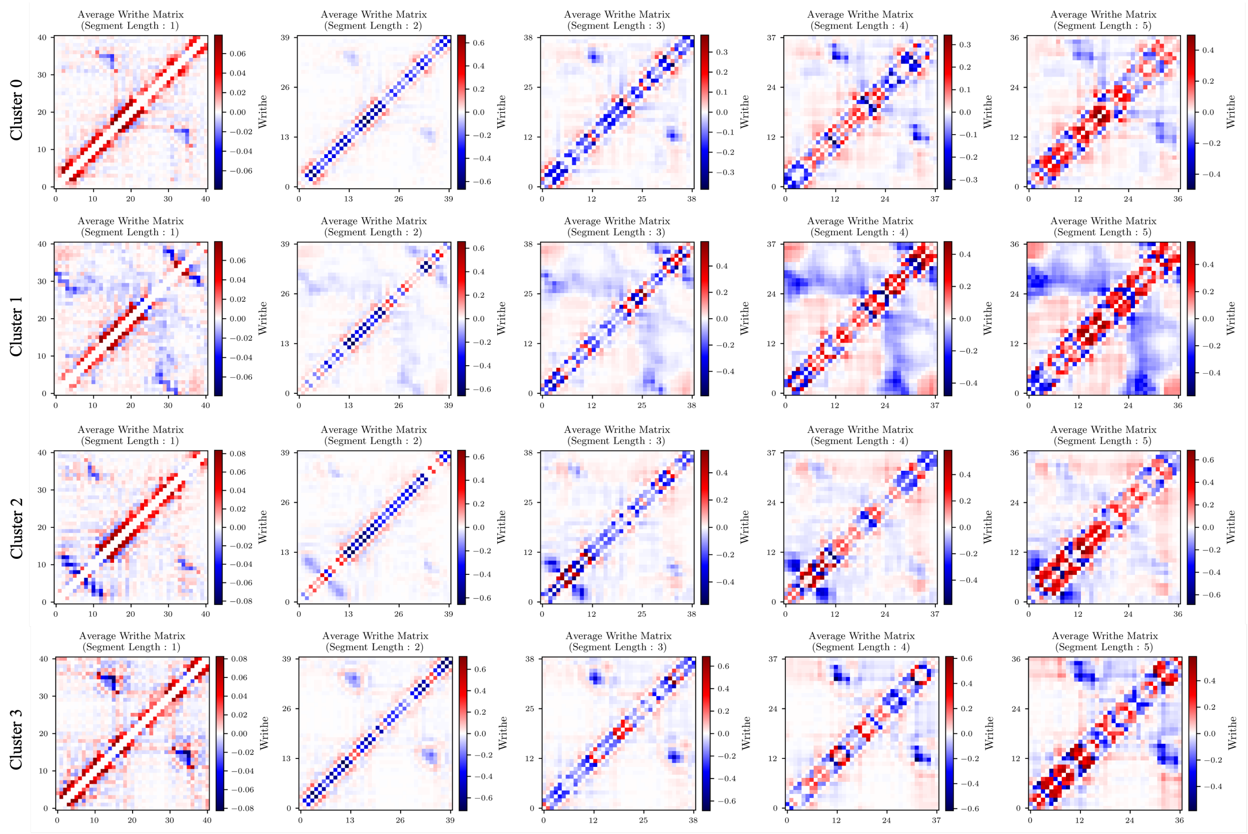


Figure S18: Pairwise writhe matrices computed within selected clusters at different segment length ($l$) ranging from 1 to 5, where the $x$- and $y$-axis indices correspond to protein residue indices of variant, $A2T_{C\beta}$.

## S7.6 A2V$_{1-6D}$

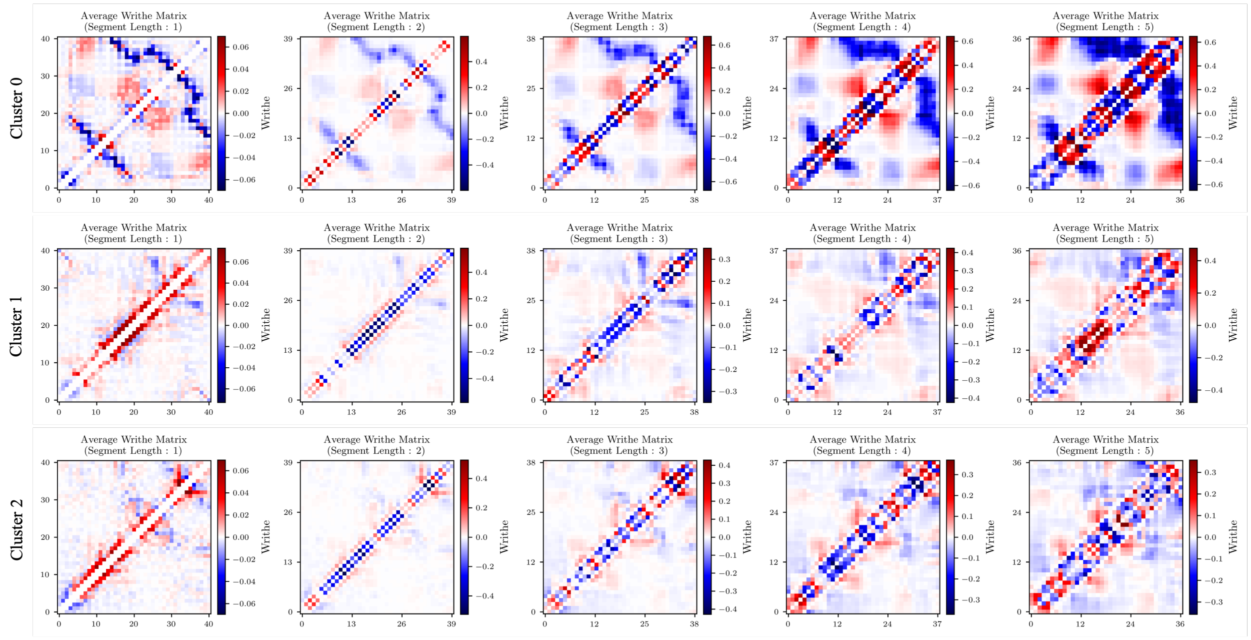


Figure S19: Pairwise writhe matrices computed within selected clusters at different segment length ($l$) ranging from 1 to 5, where the $x$- and $y$-axis indices correspond to protein residue indices of variant, A2V$_{1-6D}$.

#### S7.7 Similarity between clusters using the writhe feature as input

To define the similarity between clusters, we took the writhe feature (symmetry matrix) generated with the help of `writhe_tools`[6] and estimated their similarity using the following mathematic expression:

$$score = \sqrt{\sum_{ij} (M_{ij} - N_{ij})^2} \tag{S4}$$

where $i$ and $j$ are the indexes for two generated writhe feature matrix $\mathbf{M}$ and $\mathbf{N}$. $\mathbf{M}$ and $\mathbf{N}$ are of the same dimension. The smaller score indicates the higher similarity between two clusters.

Table S3: Summary of pairwise similarity scores computed between clusters belonging to different variants.

| | A2V | | | | A2T | | | | | $WT_{1-6D}$ | | | | | | $A2T_{C\beta}$ | | | | $A2V_{1-6D}$ | | |
|---|---|---|---|---|---|---|---|---|---|---|---|---|---|---|---|---|---|---|---|---|---|---|
| WT | 0 | 1 | 2 | 3 | 0 | 1 | 2 | 3 | 4 | 0 | 1 | 2 | 3 | 4 | 5 | 0 | 1 | 2 | 3 | 0 | 1 | 2 |
| 0 | 3.369 | 6.139 | 6.464 | 7.721 | 3.830 | 6.056 | 9.499 | 5.677 | 9.517 | 6.477 | 7.051 | 8.084 | 9.450 | 7.461 | 8.033 | 4.421 | 5.780 | 4.002 | 6.200 | 9.456 | 4.455 | 4.742 |
| 1 | 6.121 | 7.469 | 8.462 | 8.388 | 5.278 | 7.717 | 10.674 | 7.493 | 10.187 | 8.404 | 7.994 | 9.726 | 10.070 | 9.342 | 8.943 | 5.955 | 7.355 | 7.494 | 6.865 | 10.680 | 5.933 | 5.852 |
| 2 | 5.681 | 7.273 | 7.659 | 8.515 | 5.447 | 7.372 | 10.384 | 6.974 | 10.309 | 8.421 | 8.242 | 9.612 | 9.784 | 8.670 | 8.254 | 6.058 | 6.686 | 6.125 | 7.540 | 10.551 | 6.232 | 5.634 |

### S8 Secondary structure analysis of each cluster across 6 variants

## S8.1 WT

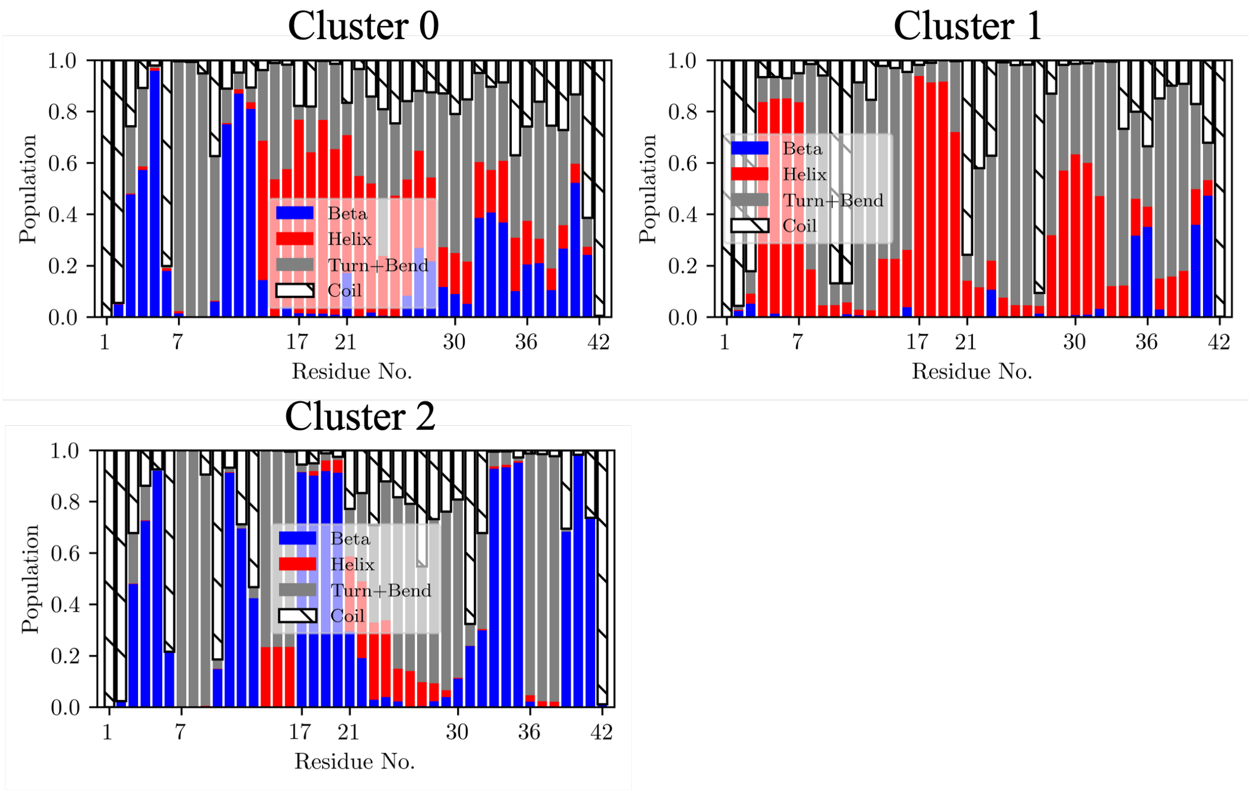


Figure S20: Per-residue secondary structure probabilities for 3 clusters of WT.

## S8.2 A2V

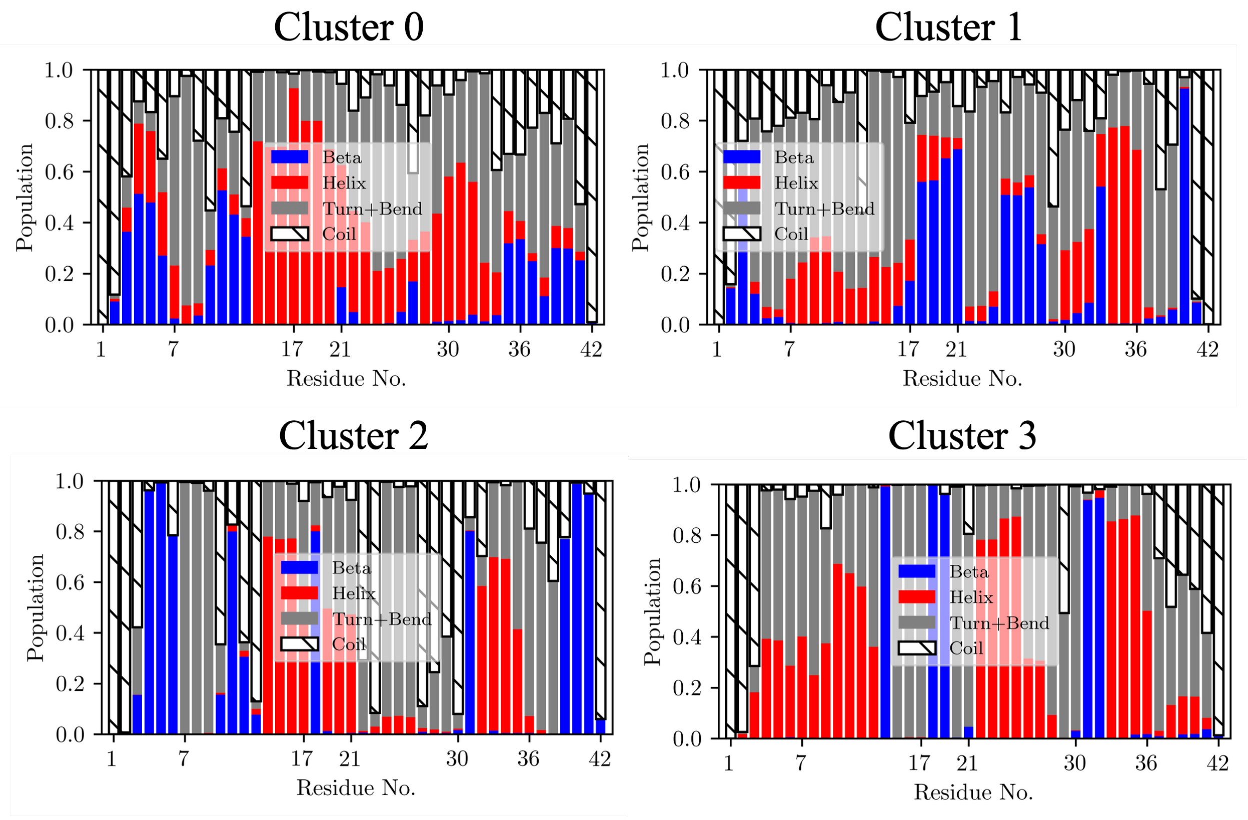


Figure S21: Per-residue secondary structure probabilities for 4 clusters of A2V.

## S8.3 A2T

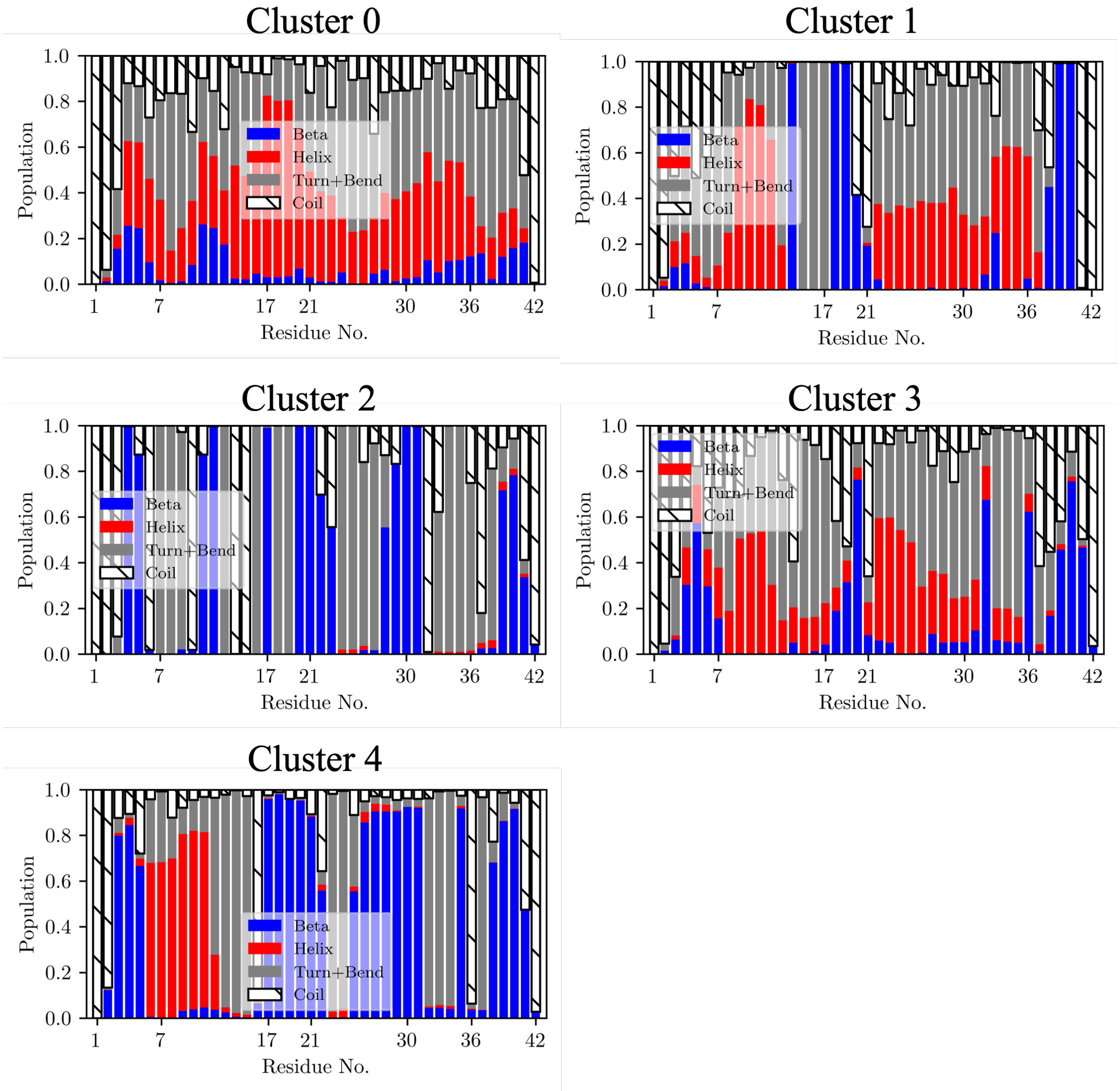


Figure S22: Per-residue secondary structure probabilities for 5 clusters of A2T.

## S8.4 $WT_{1-6D}$

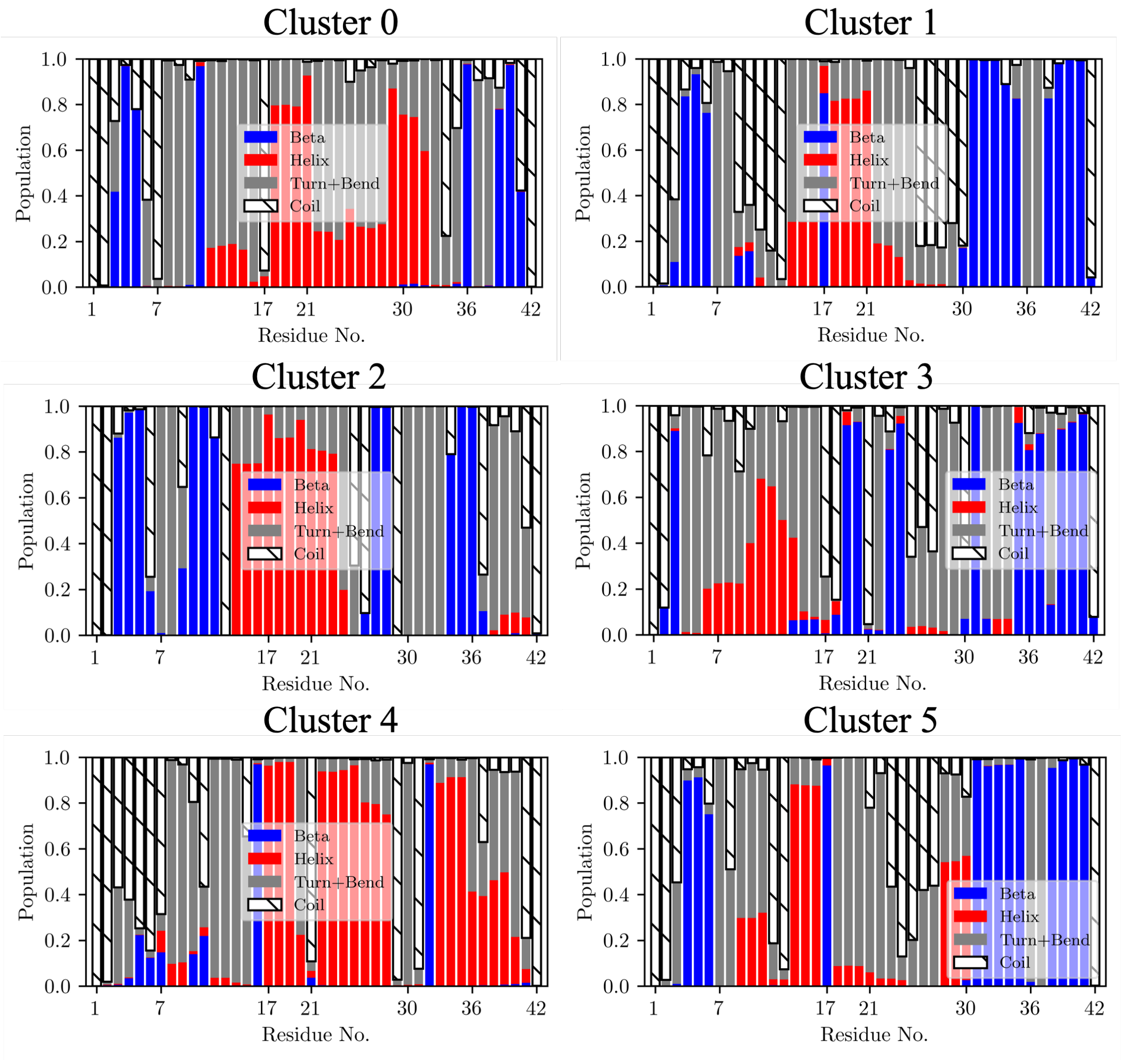


Figure S23: Per-residue secondary structure probabilities for 5 clusters of $WT_{1-6D}$.

## S8.5 A2V$_{1-6D}$

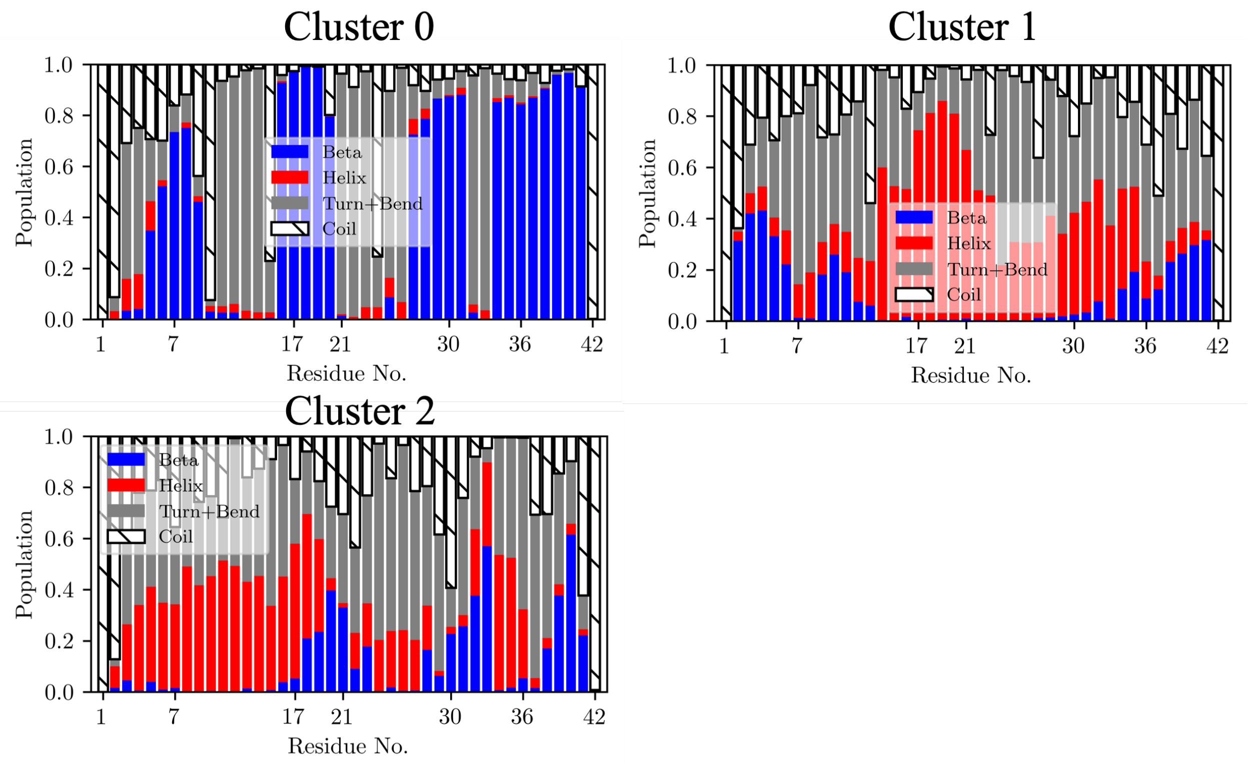


Figure S24: Per-residue secondary structure probabilities for 3 clusters of A2V$_{1-6D}$.

#### S8.6 A2T$_{C\beta}$

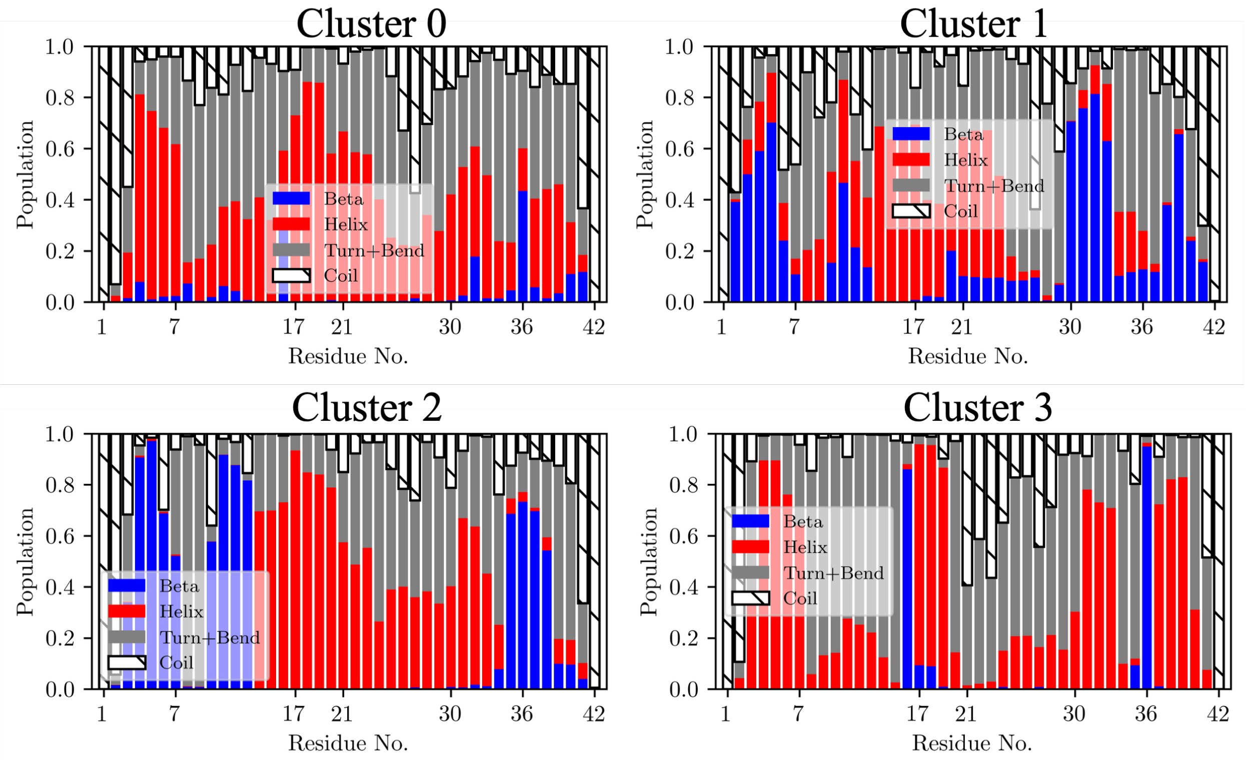


Figure S25: Per-residue secondary structure probabilities for 4 clusters of A2T$_{C\beta}$.
